# Targeted integration of extrachromosomal arrays in *C. elegans* using PhiC31 integrase

**DOI:** 10.1101/2025.11.11.687718

**Authors:** Matthew S. Rich, Ryan Pellow, Yichen Zhang, Adam Hefel, Ofer Rog, Erik M. Jorgensen

## Abstract

Extrachromosomal arrays are unique chromosome-like structures created from DNA injected into the *C. elegans* germline. However, they are unstable unless integrated into a chromosome. Current methods for integration using X-rays or CRISPR can damage DNA and exhibit low efficiency. We demonstrate that the viral integrase PhiC31, which mediates non-mutagenic recombination between short *attB* and *attP* sequences, can be used for extremely efficient and targeted integration of arrays. Arrays were integrated by PhIAT (PhiC31-mediated Integration of Arrays of Transgenes) at *attB* sites on three chromosomes, including at a fluorescent landing pad. Moreover, integrations can be inserted at any arbitrary site in the genome by simultaneously co-injecting Cas9 RNP, an *attB* repair template, and the DNA components for the array – thereby providing sites on all six chromosomes. Single injections can integrate arrays ranging in size from 1 to 18 megabases. PhIAT makes it practical to study genomes of other organisms in the nematode; one of our strains incorporates 65% of the yeast genome at a single site in the worm genome. PhIAT will accelerate a shift from unstable extrachromosomal arrays to direct integration of arrays in *C. elegans*.

## Introduction

Extrachromosomal arrays are the workhorse of *C. elegans* transgenesis. These miniature chromosomes provide a simple, rapid and efficient way to overexpress transgenes; injecting a worm gonad with a DNA mix can yield dozens of unique mini chromosomes that segregate across cell divisions and generations (*1*). Injected DNA concatenates via homologous recombination and non-homologous end-joining (*2*, *3*) into a megabase-scale circular mini-chromosome (*4*). Arrays form after fertilization, mature over the course of the first few cell divisions and are actively, but not perfectly, segregated by kinetochore proteins (*3*).

Because extrachromosomal arrays can be somatically lost and don’t follow Mendelian segregation, *C. elegans* geneticists have developed methods to integrate these giant molecules into the genome where they will be faithfully segregated. All current methods for integrating extrachromosomal arrays require DNA damage: double-stranded DNA breaks in both the genome and the array can be repaired by inserting the array into the genome. The most commonly used method to induce breaks is through ionizing irradiation (*5*, *6*). In this method, extrachromosomal arrays are selected which have a transmission rate much less than 75%, then irradiated. Since integration by irradiation is only ∼1-5% efficient, hundreds of F1 progeny are cloned and screened for 75% array transmission, indicating a heterozygous array in the genome. Those strains are then homozygosed to create a single integrated line. Irradiation breaks the genome very effectively, but randomly; arrays are integrated at random positions in the genome and must be mapped genetically. In addition, it is common for strains with arrays integrated by irradiation to have background mutations, ranging in size and severity from point mutations to chromosomal translocations. Although many associated mutation can be crossed out of the strain, because the endpoints of the integration site are often scrambled, it may be impossible to remove all background mutations.

The advent of CRISPR enabled targeted integration that is less mutagenic than irradiation. These methods create a double-stranded break in the genome (now at a single site of your choosing) (*7*, *8*). Integrants can then be identified by both positive and negative screening strategies. Targeting an endogenous gene provides a phenotypic selection (*7*). Fluorescent transgenes can be used as landing pads; integration knocks out the fluorescent protein, so integrants are isolated by loss of fluorescence (*9*). Alternatively, landing pads can be engineered with a split selectable marker for positive selection; for instance, split *unc-119(+)* rescue occurs only when an array is integrated at the landing pad (*8*). Across the board, whether using X-rays or CRISPR, the efficiency of integration is low: generally between ∼1-10% of animals after the integration process contain an integrated array in their genome. Because these methods rely on inducing breaks in both the genome and the array, they are mutagenic and the integrated array is not necessarily the same as its extrachromosomal progenitor. As such, there is still a need for an easy, high-efficiency, and non-mutagenic method for targeted integration of extrachromosomal arrays in *C. elegans*.

Phage have solved the problem of integrating their own DNA with high-fidelity into a host’s genome using serine integrases and short DNA sequences called attachment (*att*) sites (*10*). Virus genomes contain a single *attP* (“phage”) site; host genomes contain an *attB* (“bacterial”) site. A serine integrase encoded in the phage genome mediates a unidirectional recombination between the *attP* and *attB* sites, integrating the circular phage genome. This recombination converts the *attB* and *attP* sites to *attL* (“left”) and *attR* (“right”), preventing excision.

The phage PhiC31 integrase (hereon referred to as “PhiC31”) has been used widely for inserting transgenes into genomes, particularly in flies (*11*, *12*), mice (*13*), fission yeast (*14*), and human cell culture (*15*). Recently, we and others have shown that the integrase is active in the *C. elegans* germline, where it can integrate single-copy *attP*-containing transgenes and protein tags at genomic *attB* sites with high-efficiency (*16–18*). PhiC31 can also induce recombination between *attB* and *attP* sites located on homologous chromosomes as a way to target recombination (*18*). In this work, we show that PhiC31 integrates *attP*-containing extrachromosomal arrays at *attB* sites in the genome with very high efficiency. A single injection can yield a diverse collection of array sizes and compositions. We sequenced and assembled three arrays, one of them end-to-end, and found they range in length from 1.6 Mb to ∼18 Mb. We have named this method PhIAT (pronounced “fiat”) for **Ph**iC31-mediated **I**ntegration of **A**rrays of **T**ransgenes.

## Results

### Extrachromosomal arrays are integrated by PhiC31 at high efficiency

We tested whether PhiC31 could be used to integrate arrays into a single *attB* site in the *C. elegans* genome. DNA injected into a *C. elegans* gonad is assembled into large, circular extrachromosomal arrays that can contain many copies of each constituent DNA molecule. These arrays are not completely stable during cell division and are eventually lost in the strain. We reasoned that if an array contained multiple *attP* sites, PhiC31 could mediate recombination between the single *attB* site in the genome and any one of the mulitple *attP* sites on the array. The entire array would be integrated into the genome and the *attB* site would then be converted to an *attL* and *attR* site and no longer be a substrate for recombination. Even if PhiC31 recombination is very rare, the integration will be a stable component of the chromosome and maintained in that line, whereas the extrachromosomal array will continue to be lost in the population. If the integrated array carries a selectable marker, it will eventually become predominant in the population (Figure 1A).

**Figure 1.**
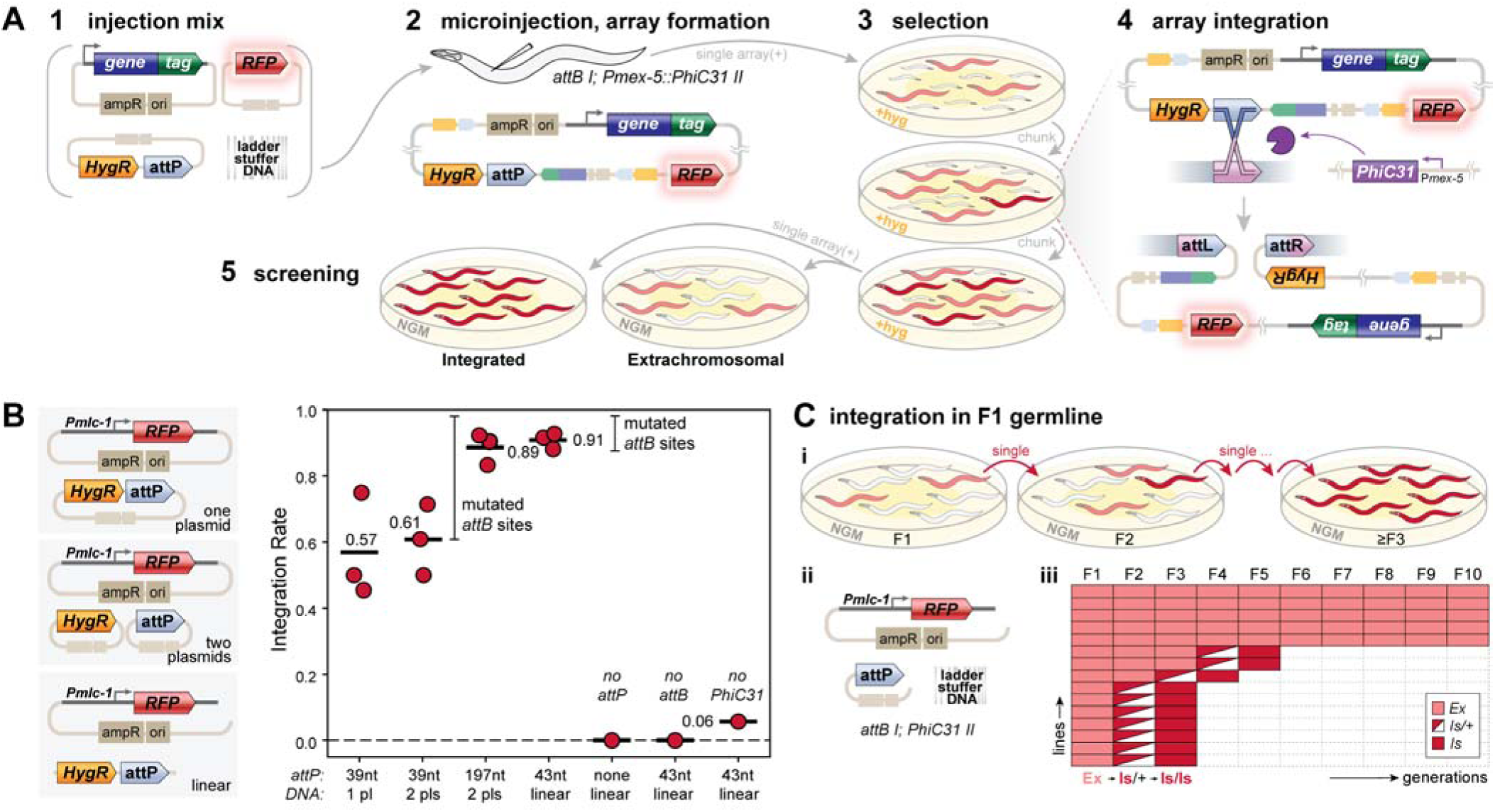
A method for targeted integration of extrachromosomal arrays using PhiC31. (A) Overview of PhIAT. (1) A DNA mixture is injected into the germline of a *C. elegans* strain that has an *attB* site in its genome (here, *oxSi1337[attB] I*) and expresses PhiC31 integrase in its germline (here, *oxSi1347[Pmex-5::PhiC31::SL2::mNeonGreen] II*). (2) Array-positive F1s are cloned to hygromycin plates and (3) grown for multiple generations while selecting for array transmission. (4) During selection, PhiC31 integrates the array into the genome. The generation after the array integrates, homozygous array-bearing animals will be in the population and out-compete non-integrated animals. (E) After selection array-bearing animals are screened for 100% transmission of the array. (B) Array integration efficiency. (left) Arrays were made by co-injecting a pan-muscle TagRFP-T transgene, a hygromycin selection (HygR), and an *attP* site using either linear DNAs, one plasmid in which the HygR and *attP* were on the same plasmid (“1 pl”), or two plasmids in which the HygR and *attP* were on different plasmids (“2 pls”). (right) Integration rates were measured for linear and plasmid-based arrays as well as controls. The type of *attP* site (197 nt, 43 nt, or 40 nt) and injected DNA types are shown on the x-axis. Each point represents, for a single injection session, the rate at which array-bearing lines that survived selection yielded a homozygous integrant. The mean rate across experiments is show as a black bar. (C) Array integration timing. (i) Outline of experiment. Array-bearing animals were cloned every generation and their progeny screened for homozygous integrants. (ii) A linear injection mix was used in this experiment. Arrays were integrated on chromosome I. (iii) A heatmap showing for each line whether the animals in that generation carried an extrachromosomal array (pink), heterozygous integrant (red/white) or an integrated array (red).

To test the method, a transgene expressing PhiC31 in the germline was inserted on chromosome II and a single *attB* site was inserted on chromosome I. These were combined to generate a strain with the *attB* site and the PhiC31 germline expression construct (hereafter ‘*attB I PhiC31 II’*). The reciprocal target *attP* site is 197 bp long, but successful recombination can also be achieved using a minimal 39 bp *attP* site in bacteria (*15*). This minimal target site was inserted into a plasmid encoding a hygromycin-resistance selectable marker (*HygR,*). The injection mix included this minimal *HygR::attP* plasmid, a pan-muscle fluorescent marker (*Pmlc-1::TagRFP-T*), and stuffer DNA; this mix was injected into the *attB I PhiC31 II* strain. Independent fluorescent array-positive F1 animals were singled and hygromycin added after 1 generation. About half of the plates did not segregate hygromycin-resistant progeny and were discarded. The plates with resistant animals were propagated on hygromycin for about 5 generations (two rounds of starvation and chunking). From each of these plates with resistant animals, 8 individuals were singled to permissive (non-drug) plates and screened for homozygous segregation of the RFP marker. Some of these animals did not segregate 100% fluorescent progeny, possibly because they were heterozygous for the integration or were extrachromosomal arrays that were still present in the population; these were discarded. Efficiency was calculated as the fraction of lines that survived selection that produced integrants: Of the 12 plates that survived selection, 6 produced homozygous inserts (50% efficiency). This experiment was repeated two more times, yielding a 61% mean integration rate of arrays made from plasmids with the minimal *attP* (Figure 1B).

Many labs include an *unc-119(+)* transgene as a selectable marker for transformation. To simplify rescue, we generated a plasmid with the minimal *attP* site and the rescuing construct for *unc-119(+)*. A strain was created containing the *attB* site, PhiC31 expression, and *unc-119*(*-*) mutation. This strain was injected with the *attP unc-119(+)* plasmid, fluorescence reporter, and stuffer DNA. The extrachromosomal array inserted into the *attB* site and rescued the Unc-119 phenotype. In addition to the HygR construct, we also created plasmids with the minimal *attP* site and the well-established drug resistance markers for neomycin/G418 (*19*) and puromycin (*20*) (Figure 2A). These plasmids were all tested for integration at the chromosome I *attB* site and drug selection successfully identified array integrations.

**Figure 2.**
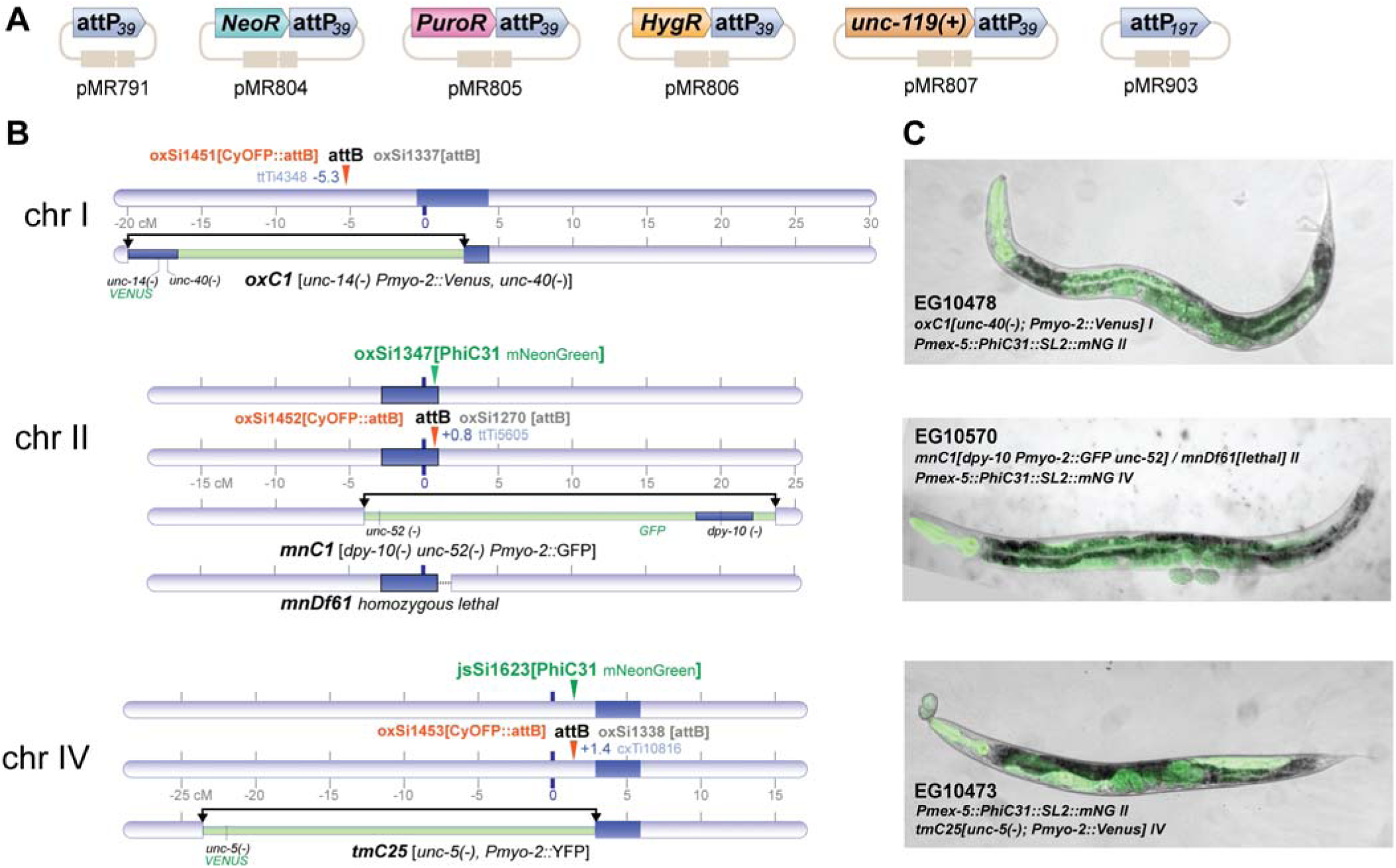
A collection of plasmids and strains for PhiC31-mediated array integration. (A) *attP* selection plasmids. pMR791 and pMR903 do not contain a selection but can be used in combination with any selection. (B) Genetic map showing locations of *attB* sites (orange), germline-expressed PhiC31 transgenes (green), and fluorescently-marked balancers with respect to chromosomes I, II, and IV. (C) To the right of each chromosome is a representative image of the PhiC31(+) balancer(+) strain for each *attB* site. Fluorescence from the balancer is visible in the pharynx of each animal. In EG10478 (top) and EG10473 (bottom), mNeonGreen fluorescence marking the PhiC31 transgene (*oxSi1347 II*) is bright.

To determine if we could improve the method by using larger *attP* recombination targets we generated *attP* plasmids with just the minimal 39 bp and 197 bp full-length sites. The HygR plasmid was injected independently, with the *attP* plasmids, the fluorescence marker, and stuffer DNA (Figure 1B). The full-length 197 bp *attP* site was significantly more efficient (89% of lines integrated) than the minimal 39 bp *attP* site (57% of lines integrated).

We then designed a slightly larger 43bp *attP* target site (*15*, *17*). Since arrays can also be assembled from injecting linear DNA, this *attP* site was appended to the HygR transgene by PCR. The resulting linear fragment was co-injected with a PCR product of the pan-muscle TagRFP-T fluorescence marker and stuffer DNA into the *attB I PhiC31 II* strain. Array-positive F1s were singled to generate lines and propagated on hygromycin plates. Arrays generated from linear DNAs integrated in 91% of lines from three sets of injections (Figure 1B).

We confirmed that integration was dependent on all three components (Figure 1B). Integration was assayed in a strain lacking the integrase PhiC31 (‘no-PhiC31’), lacking the *attB* landing pad (‘no-*attB’*), and using an array that lacked *attP* sites (‘no-*attP’*). Neither the ‘no-*attB’* nor the ‘no-*attP’* yielded integrations (out of 28 and 15 lines tested, respectively). Two arrays in the ‘no-PhiC31’ control experiment (out of 35 lines tested) yielded homozygous array insertions that did not map to the *attB* site on chromosome I.

When does integration occur? Using the above selection, integration could occur at any point before we screened for homozygous integrants. To test the timing of integration, four array-bearing animals per line were singled every generation and their progeny were screened for homozygous arrays. The generation an animal carrying a homozygous array was picked is a conservative estimate for time of integration, since integration must have occurred in the germline of an animal at least two generations prior. Integrations were tracked from 15 unique array lines over the course of 12 generations (Figure 1C). Homozygous arrays were isolated from seven of the lines in the F3 generation, implying that these arrays integrated within the germlines of the F1. Three more lines integrated by the F5 generation, but since the cloned animals could have been heterozygotes, it is possible they also represent integrations in the F1 germline. No integrations were isolated from the remaining five lines through the F10 generation.

Why do some arrays not integrate? To test whether the *attB* site could be damaged (maybe by a failed recombination) we sequenced the *attB* site from lines that survived selection without integrating an array. Out of 21 non-integrating lines, 20 had mutated *attB* sites (Figure S1A and S1B). Mutations were primarily small insertions or deletions consistent with a microhomology-dependent repair mechanism centered on the TT/AA cut site overlap (*21*, *22*).

Finally, we confirmed the specificity of integration at the *attB* site on chromosome I by genetic mapping. 38 integrants from different injections were tested, and all of them mapped to chromosome I. Altogether, these results show that PhIAT is efficient, integrates arrays specifically at *attB* sites, and happens quickly, often within the first two generations after array formation.

### Strains and DNA reagents for targeted array insertion

To facilitate use of the method in the *C. elegans* community, *attB* sites were inserted by CRISPR at safe-harbor loci on chromosomes I, II, and IV (*17*) (Figure 2B). In these strains, the *attB* sites are not marked and must be followed through crosses by PCR or by using balancers (see below). For PhiC31 transgenes, we tested our transgene on chromosome II and a similar transgene on chromosome IV (a gift from Michael Nonet). Both these transgenes are capable of high-efficiency integration (data for all integration experiments can be found in File S1).

After propagating our original strain (*attB I PhiC31 II*) for about six months, integration stopped working; sequencing demonstrated that the *attB* site had been mutated (Figure S1C). The same mutation was found six times in the non-integrating lines sequenced above. Theoretically, PhiC31 only acts on paired *attB* and *attP* sites, but at a low probability PhiC31 binding to a lone *attB* site appears to lead to cuts and mutation of the site. Typically, we inject directly into an *attB PhiC31* strain. Thus, for long-term storage, it is recommended that the *attB* site and *PhiC31* remain separated. A PhiC31-expressing strain can be crossed to an *attB* strain and homozygosed to reconstitute an injection strain.

To facilitate these crosses, both the *attB* and PhiC31 strains were altered. We added chromosomes containing a nuclear TagRFP-T transgene tightly-linked to Him mutations (*him-8 IV* and *him-5 V*) to strains containing *attB* sites. To reconstitute an injection strain, homozygous *attB* males from this strain are crossed to the PhiC31 strain. The PhiC31 strains carry a fluorescent marker that balances the *attB* site in the F1 (Figure 2B). In the F2, *attB* homozygotes will lack the balancer, PhiC31 can be homozygosed by following mNeonGreen germlines, and the Him mutation can be removed by following its linked RFP marker.

Do strains need to be homozygous for PhiC31 and *attB* to yield integrations? We tested whether heterozygous F1s generated from a reconstitution cross could yield integrations. Heterozygotes (*attB / mIn1 II; jsSi1623[PhiC31, mNG] / + IV*) were injected with pan-muscle TagRFP-T and HygR::*attP* and assayed for array integration. Two injected animals were placed on each plate and hygromycin selection was applied for two chunks of growth. 5 out of 7 lines that survived selection yielded homozygous insertions. Thus, injecting F1 heterozygotes ensures that the *attB* site is undamaged with no loss in efficiency.

### Integrated arrays are diverse

Independent arrays exhibit diverse compositions of transgenes (*1*), and individual arrays must be carefully selected for optimal function. We tested whether integrants from a single injection mix also generated a range of expression levels of a fluorescent transgene. Array-bearing F1s were singled from injected P0s. Since each oocyte in the P0 receives a different collection of DNA molecules, each F1 line is carrying a unique array. The injection mix included pan-neuronal nuclear-localized YFP (*Psnt-1::YFP::H2B*), *HygR::attP*, and yeast genomic DNA and was integrated at the *attB* site on chromosome IV. Sixteen hygromycin-resistant array-bearing lines were isolated, 10 of these lines generated integrants (63%, Figure 3A). The expression levels of YFP from these arrays ranged eight-fold from the brightest to dimmest arrays (Figure 3B). While the majority of YFP fluorescence was limited to neuronal nuclei, the brightest arrays exhibited weak fluorescence from the commissures (Figure 3C). YFP was also weakly expressed in non-neuronal nuclei in most arrays, perhaps due to leaky expression from the YFP transgene, which contained a 3’ UTR from the muscle-expressed gene *unc-54*.

**Figure 3.**
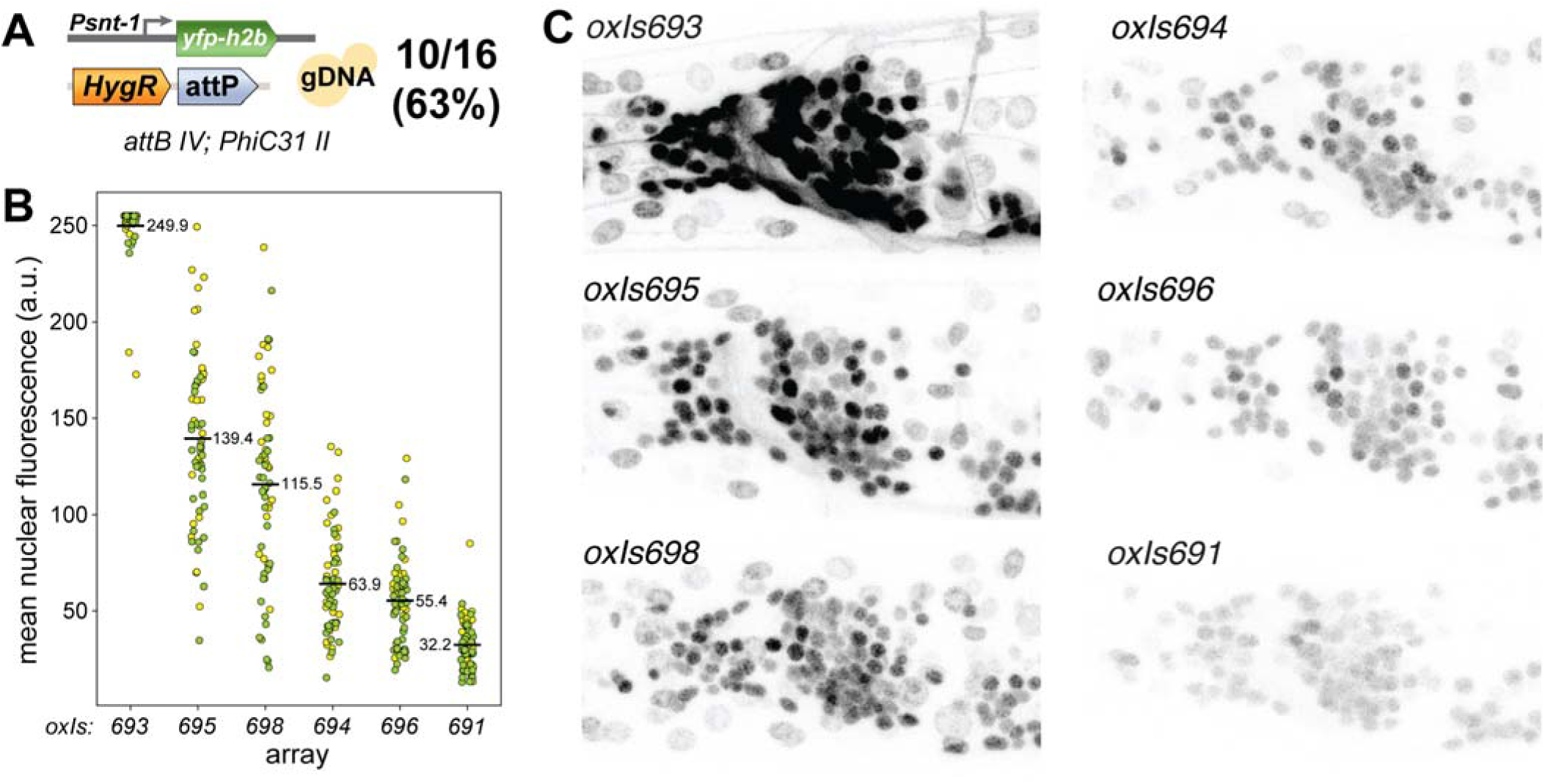
PhiC31-integrated arrays are diverse. (A) Injection mix for YFP arrays. PCR products for *Psnt-1::YFP::h2b* and *HygR::attP*, along with yeast genomic DNA for stuffer, were injected into an *PhiC31 II; attB IV* strain. 10/16 lines that survived hygromycin selection generated integrants in this experiment. (B) Animals with integrated arrays were imaged by confocal microscopy to measure the YFP brightness of their neuronal nuclei. The brightness of visible nuclei in the anterior ganglion was quantified. Each point denotes a single nucleus. Colors (yellow or green) denote measurement from two individuals. The mean fluorescence across all nuclei is annotated. (C) Representative images of integrants. YFP is largely restricted to nuclei; however fluorescence from the brightest arrays (*oxIs693* and *oxIs695*) was observed in neurites. The *oxIs693*, *oxIs695*, and *oxIs691* arrays were sequenced and assembled.

The sizes of these 10 arrays were estimated using short-read whole-genome sequencing (Table S5, see Methods for details). Seven of the 10 arrays were between 1.3 and 4.6 megabases in length, with the remaining three arrays estimated to be 7.8 Mb (*oxIs695*), 15 Mb (*oxIs698*), and 22.7 Mb (*oxIs693*). YFP brightness was roughly correlated with array size, with the exception of *oxIs698*, which was the second-largest array but near the middle of the brightness range. The amount of the yeast genome incorporated into the array was also correlated with the size of the array and ranged from 9.2% to 64.8%. Cumulatively, the ten arrays incorporated 93% of the yeast genome with 826 kb missing (Figure S2A). Reads mapped to all 16 chromosomes and the mitochondrial DNA. Fragments mapping to the yeast ribosomal DNA locus on ChrXII (XII:451,417-468,929) were present at high copy number in all ten arrays, consistent with it having a copy number of ∼150 in the yeast genome (*23*). A typical yeast gene is 1.4 kb; the mean fragment length in arrays is 12.7 kb, suggesting most of these genes are intact in the worm. To determine if there is a selection bias either for or against fragments of the yeast genome, we performed a Monte Carlo simulation. In our ten arrays, the mean amount of yeast DNA added per array is approaching, but has not reached an asymptote of 100%, suggesting that there is not a significant region of the yeast genome that cannot be incorporated into the nematode, but we cannot discount the possibility that particular genes or structural regions such as telomeres or centromeres are excluded (Figure S2B).

### Long-read assemblies of integrated arrays

The structures of extrachromosomal and integrated arrays have been characterized largely at the copy-number level by Southern blots (*1*, *2*) and imaging (*4*). Arrays are thought to be highly repetitive, which complicates assembly using short-read sequencing methods. Recently, groups have sequenced and attempted to assemble arrays using long-read nanopore sequencing (*8*, *24*, *25*), though none of these efforts were able to yield fully assembled array sequences. To improve assembly quality, we optimized an extraction protocol for isolating high molecular weight DNA for Nanopore DNA sequencing. We first used a commercial DNA ladder as stuffer DNA for the RFP arrays, which proved too short and repetitive to be assembled (*oxIs675*, not shown). Thereafter, we used yeast genomic DNA as a stuffer for the pan-neuronal YFP arrays. This DNA provided large fragments with high-sequence diversity to the array.

Three integrated arrays (*oxIs693, oxIs695, and oxIs691*) were chosen to span the range of expression levels and sizes. High molecular weight DNA from these strains was sequenced using the Oxford Nanopore MinION platform. Over 50% of Nanopore reads were longer than 15kb, and some were over 150kb in length. These lengths provided sufficient coverage to use only reads longer than 50kb for assembly (Table S6 Sequence Statistics). Contigs were further refined by manual assembly using unique polymorphisms within the array as landmarks (for example, some *Psnt-1::YFP* repeats were fragmented).

The lengths were concordant with the short-read estimates above (*oxIs691*: 1.6 Mb by short reads and 1.6 Mb assembled; *oxIs693*: 22.7 Mb by short reads and 17.9 Mb assembled; *oxIs695*: 7.8 Mb by short reads and 6.6 Mb assembled). *attL* and *attR* sites were present and undamaged at the ends of the insertion site. Annotated gene models of each array appear in Figure 4, and the details are described here:

**Figure 4.**
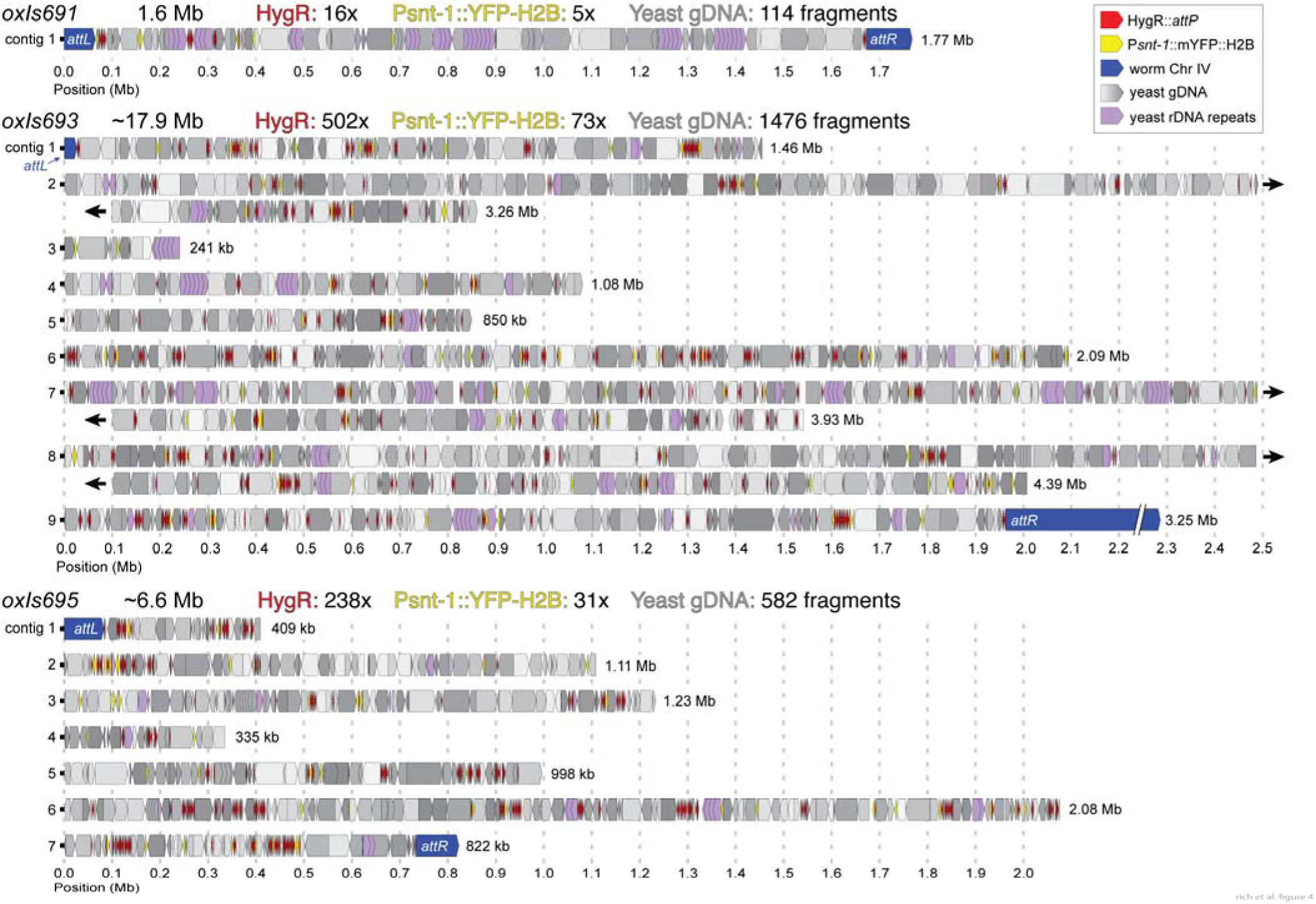
Assembled integrated arrays. Annotated contigs for *oxIs691, oxIs693, and oxIs695*. The estimated total length of each array as well as number of constituent fragments is written above each diagram. A single contig is drawn per line. The length of each contig is written at the end of each line. The positions of sequence-verified flanking *attL* and *attR* sites are noted. In *oxIs693* contigs 2, 7, and 8, were too long to plot easily and were wrapped to new lines; these cases are denoted with black arrowheads. An additional megabase of *C. elegans* sequence has been truncated from the right side contig 9 in *oxIs693* for the purpose of drawing this figure. Annotations are drawn as boxes with arrows pointing in the direction of transcription (for transgenes) or based on the reference (for yeast or worm genome fragments) and are colored based on their identity (HygR: red; YFP: yellow, *C. elegans* chromosome IV: blue; yeast genomic DNA: gray, shaded based on chromosome; the yeast rDNA repeat from chromosome XII: purple).

#### oxIs691

*oxIs691* is the dimmest array of the set and is also the smallest at 1.6 Mb long. We obtained ∼279K reads longer than 10 kb and assembled the array with no gaps including the expected flanking sequences to the *attL* and *attR* sites on chromosome IV. The array contains 5 full-length YFP fragments (out of 6 total, 83%), 16 full-length HygR fragments (out of 24 total, 67%), and 111 yeast fragments (Figure S3A-C). Yeast gDNA fragments ranged from 76 bp to 65,673 bp in length, with 91.5% longer than 1000 bp. 2.3% of the array was made of HygR fragments, 1.6% was made of YFP fragments, and 96.1% was the yeast genome, largely reflecting the molar ratios of injected DNA.

#### oxIs695

This is the second-brightest array. *oxIs695* is ∼6.8 Mb long and was integrated in the correct location on chromosome IV flanked by *attL* and *attR* sites. We obtained 185K reads longer than 10 kb and were able to assemble this array into seven contigs ranging from 335 kb to 2.1 Mb (N50=998 kb). The array contains 32 full-length YFP transgenes (from 54 total fragments, 59%), 238 full-length HygR transgenes (out of 384 total fragments, 62%) and 558 yeast fragments. Yeast DNA fragments ranged from 45 bp to 74,829 bp in length; 86% were longer 1000 bp. Yeast genome comprised 88.9% of the array, 8.1% was made of HygR fragments, 3% was made of YFP fragments. Fragment lengths can be found in Figure S3D-F.

#### oxIs693

This is the brightest array. *oxIs693* is ∼17.9 Mb long and was integrated in the correct location on chromosome IV flanked by *attL* and *attR* sites. We were able to assemble this array into nine contigs ranging from 240kb to 4.4Mb (N50=2.3 Mb). The array contains 73 full-length YFP fragments (out of 109 total, 67%), 502 full-length HygR fragments (out of 772 total, 52%), and 1446 yeast fragments (Figure S3G-I). Yeast gDNA fragments ranged from 43 bp to 85,495 bp in length; 83% were longer 1000 bp. Yeast genome comprised 92% of the array, 5.8% of the array was made of HygR fragments, 2.2% was made of YFP fragments.

We were unable to fully assemble the two largest arrays *oxIs693* and *oxIs695* due to the presence of large duplications within the array. Each array has 21 regions longer than 50 kb that are duplicated in multiple contigs (Figure S4). These repeats are composed of multiple DNA fragments, implying that domains of linear DNA became concatenated first, then either duplicated and merged by recombination, or were breaks in the array that were repaired by replication.

During formation of extrachromosomal arrays, injected circular DNAs recombine into tandem arrays, whereas linear DNAs assemble randomly by end-joining (*8*, *24*, *25*). At first glance, our arrays made from linear DNA are consistent with random assortment, but the HygR fragments appear to be clustered. Fragment clustering was tested using a Monte Carlo simulation: DNA fragments in the arrays were randomly assorted and then scored for whether neighbors were the same fragment type (yeast, HygR, or YFP). The HygR transgenes in our arrays were significantly clustered (p<1e-5 in both arrays), whereas YFP transgenes in the array were randomly assorted within the array (p=0.16 in *oxIs695* and p=0.42 in *oxIs693*). It is not yet clear why HygR genes would be clustered. It is possible that concatenation of these fragments is a property of the DNA; alternatively clustering is due to selection of arrays for high expression of the HygR gene.

### One-shot attB and array integration

The strains constructed for this manuscript contain *attB* sites near common landing pads on chromosomes I, II, and IV. *attB* sites are short and simple to edit into the genome by injecting Cas9 ribonucleoprotein complex and a single-stranded DNA repair donor (*2*). We tested whether we could integrate an array at an arbitrary site in the genome through two sequential steps from a single injection mix: (1) an *attB* site is edited into the genome using CRISPR reagents, probably taking place in the germline of the P0 or the F1 zygote, then (2) the array that formed from co-injected DNA is integrated into the *attB* site by PhiC31, which probably occurs in the F1 germline (Figure 5A).

**Figure 5.**
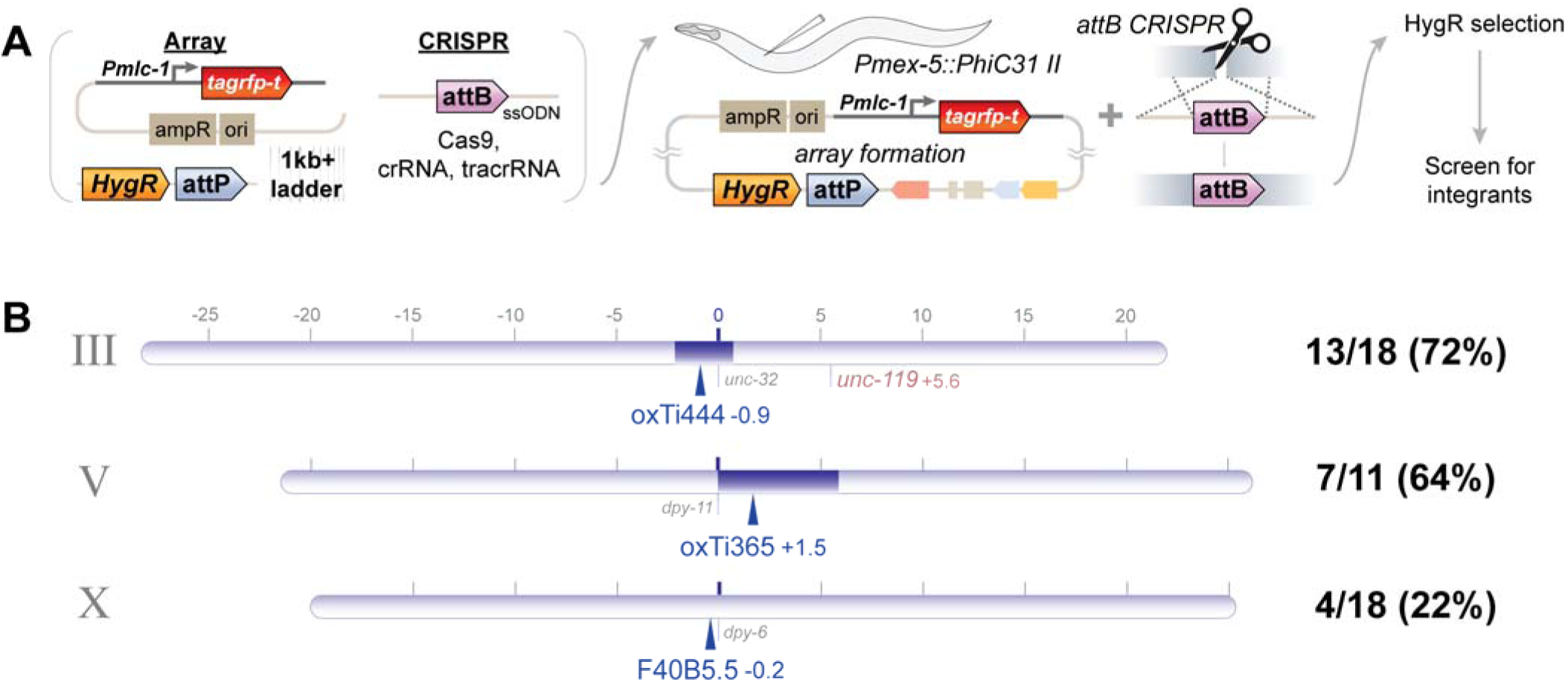
One-shot PhIAT: attB knock-in and array integration from a single injection. (A) Overview of method. A PhiC31(+) strain is injected with a single mix containing DNA reagents for an array and CRISPR reagents for knocking an *attB* site into the genome with and ssODN and Cas9 ribonucleoprotein complex. Array-bearing F1 animals, which have also likely received CRISPR reagents that can insert an attB site in the genome, are cloned, selected, and screened for integrants. (B) Genetic map for three sites in the genome using ‘one-shot PhIAT’: near the *oxTi444* locus on chromosome III, near the *oxTi365* locus on chromosome V, and near *F40B5.5* on chromosome X. The frequency at which we isolated integrants at each site is shown to the right of the chromosome.

A PhiC31-expressing strain was injected with the CRISPR reagents to insert an *attB* site on chromosome III. The injection mix also contained PCR products for pan-muscle TagRFP-T and HygR::*attP*. F1 array-bearing animals were cloned and selected. Of the 18 lines that survived selection, 13 yielded a homozygous array integration (72%). Genetic mapping of these insertions placed them on chromosome III consistent with being inserted at the new *attB* site. The same experiment was repeated targeting sites on chromosomes V and X, yielding 7 integrants from 11 lines with an insertion for chromosome V (64%) and 4 integrants from 18 lines for chromosome X (22%) (Figure 5B).

To determine whether these arrays were integrated by PhIAT, short-read shotgun sequencing was used to characterize the insertion breakpoints. An array inserted by PhIAT would have *attL* and *attR* scars between the *C. elegans* genomic sequence and the array. Out of 17 lines sequenced, 11 (65%) had a correct *attR* site at the junction with the chromosome; apparently an *attB* site had repaired into the cut site, followed by array integration. Four of the lines did not have *attR* sites at the junctions. These arrays had inserted into the Cas9 cut site using a repair mechanism similar to FLInt (*9*) before the *attB* site had healed the double-strand break. The junction could not be identified in two of the lines.

Altogether, these experiments show that arrays can be inserted at new *attB* sites using a single injection, albeit with lower efficiency than injecting into a strain with an existing *attB* site. These arrays are integrated predominantly by an initial *attB* knock-in followed by PhiC31-mediated recombination. Thus, efficient array integration can be achieved anywhere there is an efficient Cas9 cut site.

### Integration at a fluorescent landing pad

The three *attB* landing pads on chromosome I, II, and IV are unmarked, simply because short *attB* sites are easy to knock into the genome. Integrations into these sites are detected by visually assaying for homozygous transmission. An alternative method for inserting arrays is to use Cas9 to cut a fluorescent transgene inserted at an existing miniMos insertion using FLInt (*9*, *26*). Thus, FLInt provides both a positive screen (presence of the array) and negative screen (loss of landing pad fluorescence). To implement this strategy using PhIAT, an *attB* landing pad was inserted in an intron of a CyOFP transgene expressed in the intestine (Figure 6). A single copy of this landing pad was integrated on chromosomes I, II, and IV. The landing pad on chromosome I was crossed to a strain expressing PhiC31. The strain was injected with linear DNAs encoding a pan-muscle TagRFP-T transgene and a HygR::*attP* PCR product. Array-bearing lines were selected on hygromycin as before and examined for loss of CyOFP expression in the intestine. In 11 of 25 F1 lines, hygromycin-resistant integration was associated with loss of CyOFP expression (44% integration rate). One additional line integrated a hygromycin-resistant array but did not lose CyOFP fluorescence. Another line lost CyOFP fluorescence but did not integrate its array. These experiments demonstrate that array integration can be identified based on disruption of a fluorescent landing pad at rates similar to a simple *attB* site in the genome.

**Figure 6.**
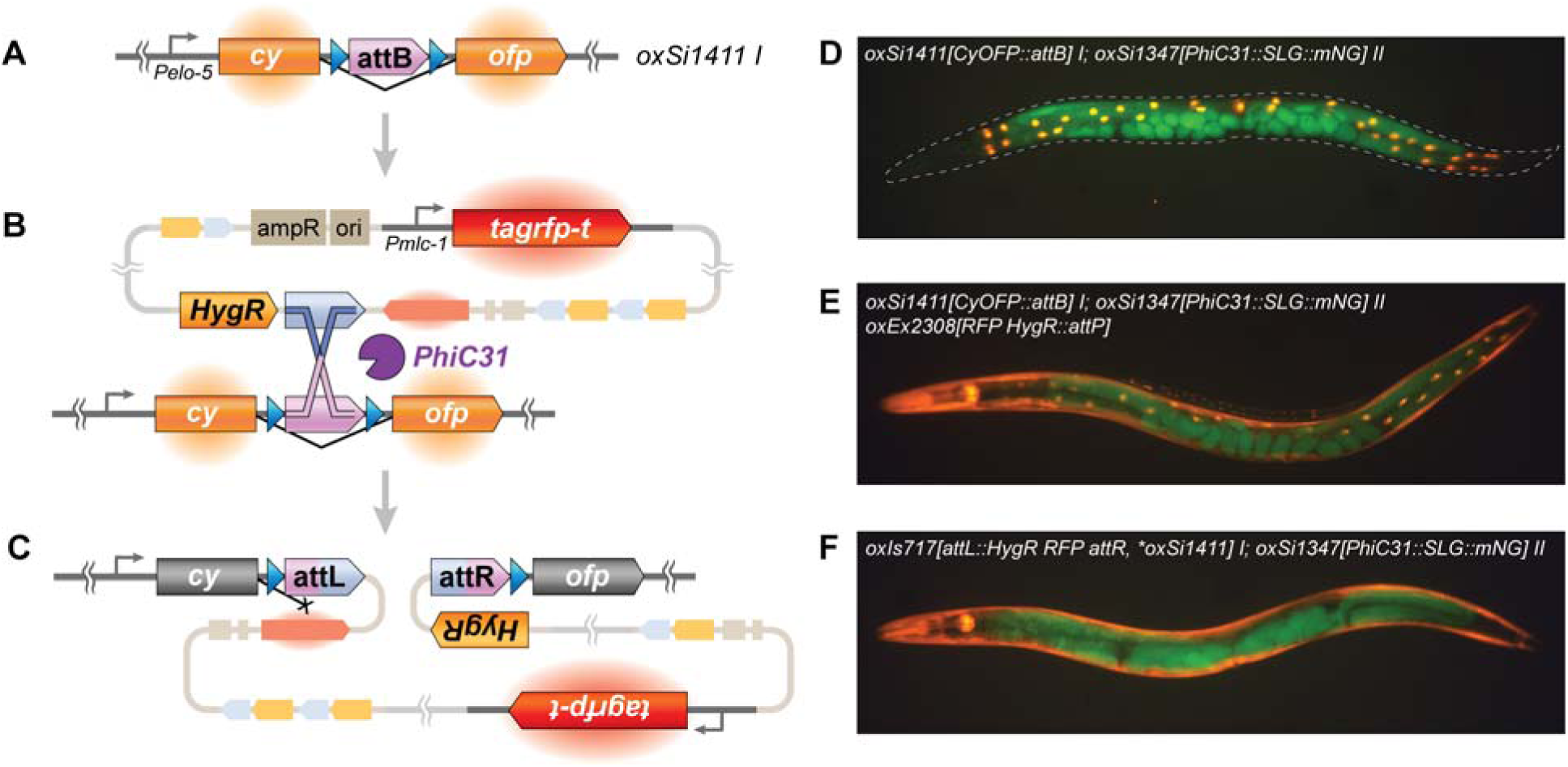
Integration into a fluorescent landing pad. (A-C) Schematic of integration into a landing pad. (A) An *attB* landing pad was inserted in an intron of a single-copy transgene expressing a nuclear-localized CyOFP in the intestine on chromosome I. (Note the *attB* site is flanked by *loxP* sites (blue), which were included to test whether the array could be excised after integration; these experiments were not successful). (B) An extrachromosomal array (in this case, containing muscle-expressed TagRFP-T, *attP* sites, and hygromycin selection) is injected into the landing pad strain. During selection, PhiC31 mediates recombination between *attP* in the array and *attB* in the genome. (C) When the array is integrated, it is recombined into the CyOFP intron, knocking out the fluorescence. Fluorescence from the array is maintained. (D-F) Representative images of animals at each stage in the integration process. In (D), CyOFP(+) intestinal nuclei can be seen. In (E), the strain carries an extrachromosomal array before integration; both CyOFP(+) intestinal nuclei and TagRFP-T fluorescence from the array can be seen. In (F), the array from (E) has integrated into the landing pad and is homozygous; no CyOFP fluorescence is present in the intestine. Germline mNeonGreen fluorescence from the PhiC31 transgene is present in all images.

### Stability and consequences of large integrations

PhIAT integrants contain a high number of repeats from the injected DNA, and can be very large, in some cases nearly doubling the size of the chromosome. For example, chromosome IV is 17.5 Mb; our longest integrant adds 18 Mb (*oxIs693*). Large repetitive insertions could be unstable due to recombination within the array or interfere with chromosome segregation due to their massive size. We tested for chromosome instability using three assays.

First, array stability was tested by loss of fluorescence. Eighteen independent integrants were chosen. These strains had arrays inserted at either the chromosome I or IV landing pads, and carried a hygromycin-resistance transgene, a fluorescent marker, and either DNA ladder or complex yeast DNA as stuffer. A cohort of eggs were collected overnight (∼60-120 embryos each); these progeny were each examined for reduced fluorescence. All progeny from each of the 18 integrants exhibited no changes in brightness (Table S7).

Second, some of these arrays are the size of chromosomes, and could possibly interfere with segregation of autosomes. In particular, three arrays are larger than 7 Mb. If the presence of an integrated array interferes with the normal pairing of the chromosome, then the homozygous array, and perhaps especially an unpaired heterozygous array, would disrupt chromosome segregation in meiosis and lead to reduced fecundity or even sterility. Seven independent integrations on chromosomes I and IV were chosen from the set above (Table S7) and tested as homozygotes or crossed to the wild type (N2) and tested as heterozygotes. The seven integrants had largely similar progeny counts as either homozygotes or heterozygotes (means ranging from 200 to 278 progeny, Figure S5A). In five of the seven integrants, one of the tested lines was an outlier with low fecundity (2 homozygotes, 3 heterozygotes). To test whether this was heritable (and potentially linked to the array), the fecundity of their offspring was retested (Figure S5B). In all five cases, normal fecundity was observed, though one line from retesting *oxIs695* was sterile.

Third, in *C. elegans* males, the single X chromosome is unpaired (genotype XO, in which ‘O’ is designated as the absence of a second X chromosome). Segregation of the X chromosome from the integration site would lead to sperm carrying the integrant but lacking the X chromosome. Previous work has demonstrated males carrying integrated arrays produce an excess of hermaphrodites lacking the array and male progeny that carry the array (*27*). We tested nine of our arrays by crossing males heterozygous for the array to wild-type hermaphrodites. The fraction of array transmission to sons and daughters was calculated. Seven of the nine arrays segregated from the X (male frequency > 0.67, Table S7). The three assembled arrays with precisely known lengths were among those tested; *oxIs691* (IV 1.7Mb) did not segregate from the X chromosome (0.43), whereas *oxIs693* (IV ∼17Mb) and oxIs695 (IV ∼8Mb) exhibited strong segregation from the X chromosome (0.96 and 0.87, respectively).

On a practical level, the integrant does not affect X-chromosome segregation in hermaphrodites; the two X-chromosome pair and segregate from each other normally. However, when crossing array-bearing males into different genetic backgrounds, obtaining hermaphrodites with the array will be more difficult. In the most extreme case, the resulting offspring resulted in 96% of array-positive animals being males, and only 4% were hermaphrodites.

## Discussion

For extrachromosomal arrays to be maintained with stable expression and segregation, they must be integrated into the genome. PhIAT is a method for high-efficiency targeted integration of extrachromosomal arrays in *C. elegans*. Arrays are generated by injecting DNA along with PhiC31 *attP* sites, these are then integrated into *attB* sites in the genome the next generation. If the array carries a selectable marker, the integration eventually outruns the unstable extrachromosomal array. After ∼5 generations, the integration is recovered with ∼50-100% efficiency among surviving animals. The most important benefits of the method are that the insertion is at a known site in the genome, integration is very efficient, insertion by recombination is non-mutagenic, and a single injection can yield many highly diverse arrays.

In comparison, the best current methods for array integration at a targeted site use Cas9 / CRISPR to cut the genome at a specific site, namely mosTI (*8*) and FLInt (*9*). The published efficiencies of mosTI and FLInt are lower than PhiAT, but not directly comparable because they represent measurements after a single generation and will consequently appear lower. But under hygromycin selection as described here, integrants using mosTi and FLInt should also outrun extrachromosomal arrays – the tortoise always wins – and probably exhibit higher efficiencies. Another potential consideration is that double-strand breaks can lead to broader DNA damage. These can be ameliorated in mosTI and FLInt by outcrossing, but they remain a concern.

The most important concern using PhiAT is that the PhiC31 recombinase mutates *attB* sites in the absence of an *attP* target. This precludes the long-term maintenance of an injection strain that constitutively expresses PhiC31 and carries an *attB* landing pad. However, the injection strain can be easily generated by crossing males bearing the landing pad (*attB Him*) to the PhiC31 expressing strain. In the future, inducible germline expression would improve the convenience of PhIAT. Heat-shock can drive the expression of genome-modifying transgenes like Mos1 transposase (*28*), Cas9 (*29*), and Cre recombinase (*30*), but expression is only transient. The bipartite TetON system works well in somatic cells (*18*, *31*), but our attempts to induce expression in the germline have been unsuccessful (unpublished). Finally, injection of purified PhiC31 enzyme (*32*) could be used to integrate existing extrachromosomal arrays into the attB landing pad in the same way that Cas9 can be delivered by injection for CRISPR (*33*).

An important benefit of PhIAT is the number and diversity of integrants one can isolate. Here, a single researcher isolated a total of 203 integrated arrays across 25 injection sessions, including one session that yielded 24 unique integrants. Such integrants incorporate diverse ratios and amounts of injected DNA. Array diversity is important for obtaining the correct stoichiometry of transgenes. For example, the composition of the NeuroPAL array, used for identifying neurons by color and position, needed to be tuned to optimize expression of four different fluorescent proteins among 41 transgenes (*34*). It is feasible to create up to a hundred stably integrated arrays by PhIAT to isolate an optimal array based on function.

Our assemblies supplement the small but growing set of sequenced arrays in *C. elegans* (*8*, *24*, *25*). Ten arrays were sequenced using short-read methods, three of these were further sequenced by long-read Nanopore. One of them, *oxIs691,* is a complete 1.6Mb end-to-end assembly. These reconstructions provide a resource for the study of how extrachromosomal DNA assembles in *C elegans*. For the most part the injected linear fragments assembled randomly. Two anomalies were observed; first, some arrays contained complex assemblages that were duplicated, second, the HygR gene repeats were non-randomly clustered. Whether these anomalies form during extrachromosomal array formation, array integration, or are the result of selection is not known.

Despite the presence of large arrays of containing repeats, integrated arrays appear to be stable during propagation. Two of these arrays are enormous: ∼7 Mb (*oxIs695*) and ∼18 Mb (*oxIs693*). These two strains are noticeably sick and grow slowly, possibly due to overexpression of transgenes in the array, or alternatively due to broader effects on the endogenous genome. The presence of the large insertion as homozygotes or heterozygotes does not appear to affect segregation of chromosomes in mitosis or meiosis in hermaphrodites. Moreover, *C. elegans* can tolerate large chromosomes: strains carrying end-to-end chromosome fusions (*mnT12 X;IV* or *meT7 III;X;IV*) are phenotypically wild-type (*35*, *36*). However, large insertions have a profound effect during male meiosis; the array-bearing chromosomes segregate from the X-chromosome in XO males. ‘Distributive pairing’ and segregation has been well described in flies. Segregation does not depend on sequence identity, crossovers, or even chromosome pairing, but rather simply on chromosome size and limited real estate of the meiotic spindle (*37*, *38*).

Finally, we note that these arrays are largely composed of yeast DNA. The yeast genome is 12.1 Mb; sequence data from 10 arrays accounts for 93% of the yeast genome. One single array covers 63% of the yeast genome, begging the question why the strain is still nematode and not yeast. These methods in the future could allow *C. elegans* to be used as a crucible for studying the evolution and function of genomes of other organisms, transgressing boundaries between species.

## Supporting information

Supplemental File 3 (Detailed Protocol)

## Acknowledgments.

Michael Nonet and Christian Frokjaer-Jensen provided important insight in the development of the method. We also thank Michael Nonet for sharing unpublished reagents as well as results regarding *attB* mutation. We thank Maitreya Dunham for reminding us where the yeast ribosomal DNA locus was. Markell Kolendrianos helped with worm maintenance during selections. We thank the rest of the Jorgensen Lab for helpful suggestions for both experiments and writing. Genome sequencing was performed at the Huntsman Cancer Institute High-Throughput Genomics core. We thank the *Caenorhabditis* Genome Center (University of Minnesota) for strains and Wormbase and the *Saccharomyces* Genome Database for maintaining genomic and genetic data.

## Author Contributions

MSR and EMJ conceived of the project. MSR and AH created *attB* and *PhiC31* strains. Long-read sequencing and analysis was performed by RP and MSR. MSR and YZ performed genetics experiments. MSR and EMJ wrote the manuscript. EMJ and OR acquired funding.

## Data, Code, and Materials Availability

All data and code needed to evaluate the conclusions in the paper are present in the paper and/or the Supplementary Materials. Raw sequencing reads can be found in NCBI SRA Bioproject PRJNA1493141. *C. elegans* strains created in the manuscript are deposited at the Caenorhabditis Genetics Center (University of Minnesota). Plasmids created in the manuscript are deposited at Addgene.

---

**Figure S1.**
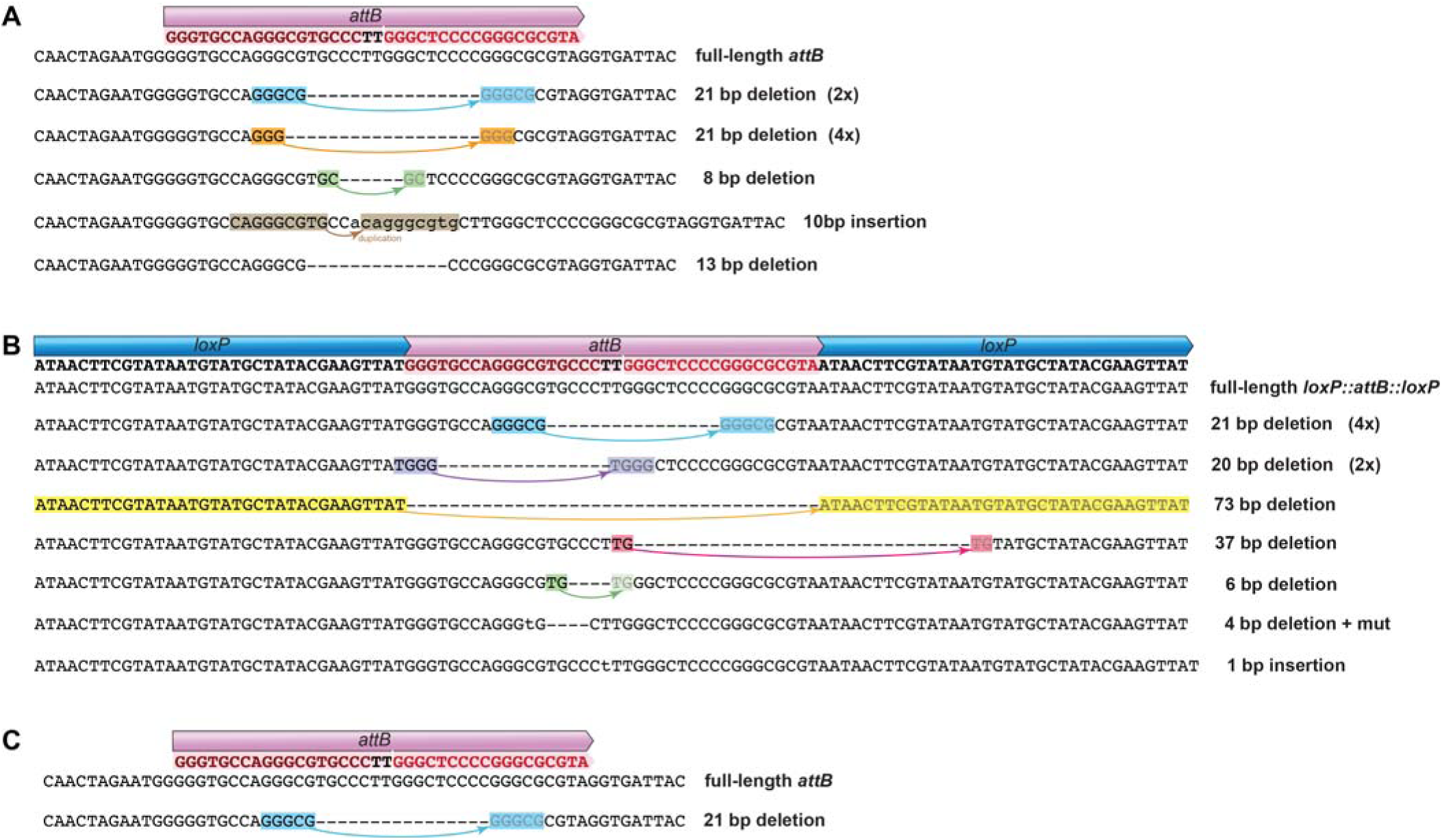
Mutated *attB* sites. We sequenced the *attB* sites in strains that did not yield integrants. Each line is a different mutation to the *attB* site, with a description and whether it was isolated multiple times shown to the right. Most mutations were consistent with a microhomology-mediated repair causing a deletion. These repairs are annotated with colored highlights showing the relevant microhomology in which gray residues are now absent. (A) Mutated *attB* sites from attempted integrations into *oxSi1337*, (B) into *oxSi1411*. (C) The mutation found in *oxSi1337* after long-term propagation with PhiC31. This mutation was also found in (A) and (B).

**Figure S2.**
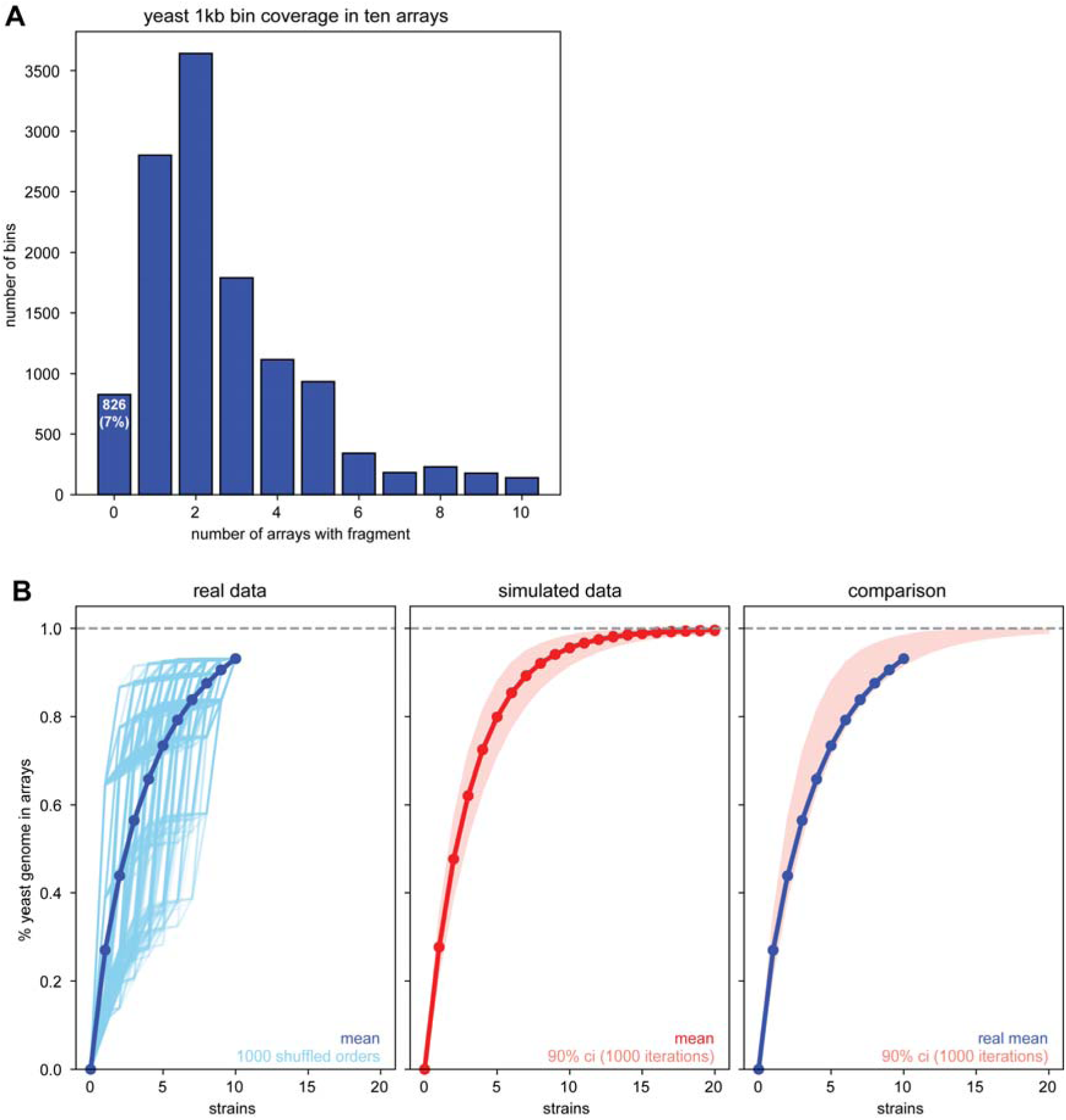
Yeast fragments in integrated arrays. (A) Number of times yeast genomic fragments are found in 10 sequenced arrays. The yeast genome was split into 1000 bp bins, and whether the sequence in each bin was present in an array was determined by short-read whole genome sequencing. 93% of the yeast genome is found in at least one of the ten arrays. (B) Saturation of yeast genome in 10 arrays. (Left) Average saturation (dark blue) was calculated using a Monte Carlo simulation shuffling the order that arrays were added to the cumulative plot (light blue lines). For example, the left-most light blue line is the order in which arrays containing the most yeast DNA were the sampled first, and the right-most line is the order in which the arrays containing the least yeast DNA were sampled first. (Middle) An expected distribution for incorporation of yeast fragments into arrays was created using a Monte Carlo simulation (see Methods). (Right) Overlay of the real mean saturation with the 90% confidence interval from simulated data. The real arrays (blue line) grow slower than expected (but within the confidence interval). This is likely due to the yeast rDNA being overrepresented in the injected DNA (∼150 copies) compared to the reference genome (2 copies).

**Figure S3.**
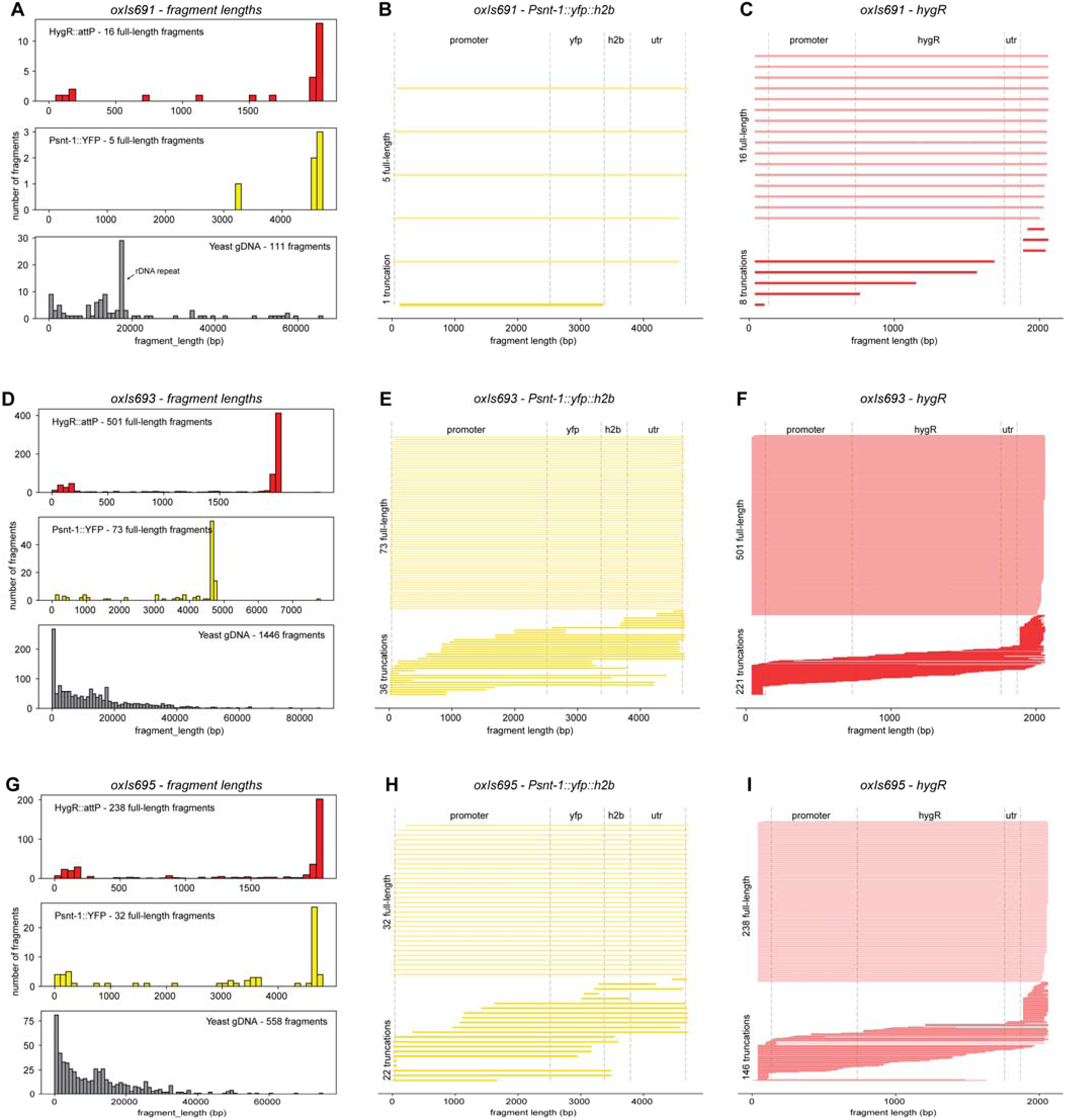
Transgene fragments in assembled integrated arrays. Contigs from *oxIs691*, *oxIs693*, and *oxIs695* were aligned to the yeast genome, P*snt-1::YFP*, and *HygR::attP* references. Mapped fragments were filtered for those having a mapping quality > 0. For *HygR::attP* and P*snt-1::YFP*, being called as a “full-length” fragment required a mapping length of at least 95% of the reference length. For *Psnt-1::YFP* and *HygR* fragment plots, each mapped fragment is a horizontal line, with full-length fragments on top and truncated fragments on the bottom of the plot. Boundaries between functional regions are shown by vertical dashed lines. *oxIs691:* (A) fragment length distributions; (B) *Psnt-1::YFP* fragments; (C) *HygR* fragments. *oxIs693:* (D) fragment length distributions; (E) *Psnt-1::YFP* fragments; (F) *HygR* fragments. *oxIs695:* (G) fragment length distributions; (H) *Psnt-1::YFP* fragments; (I) *HygR* fragments.

**Figure S4.**
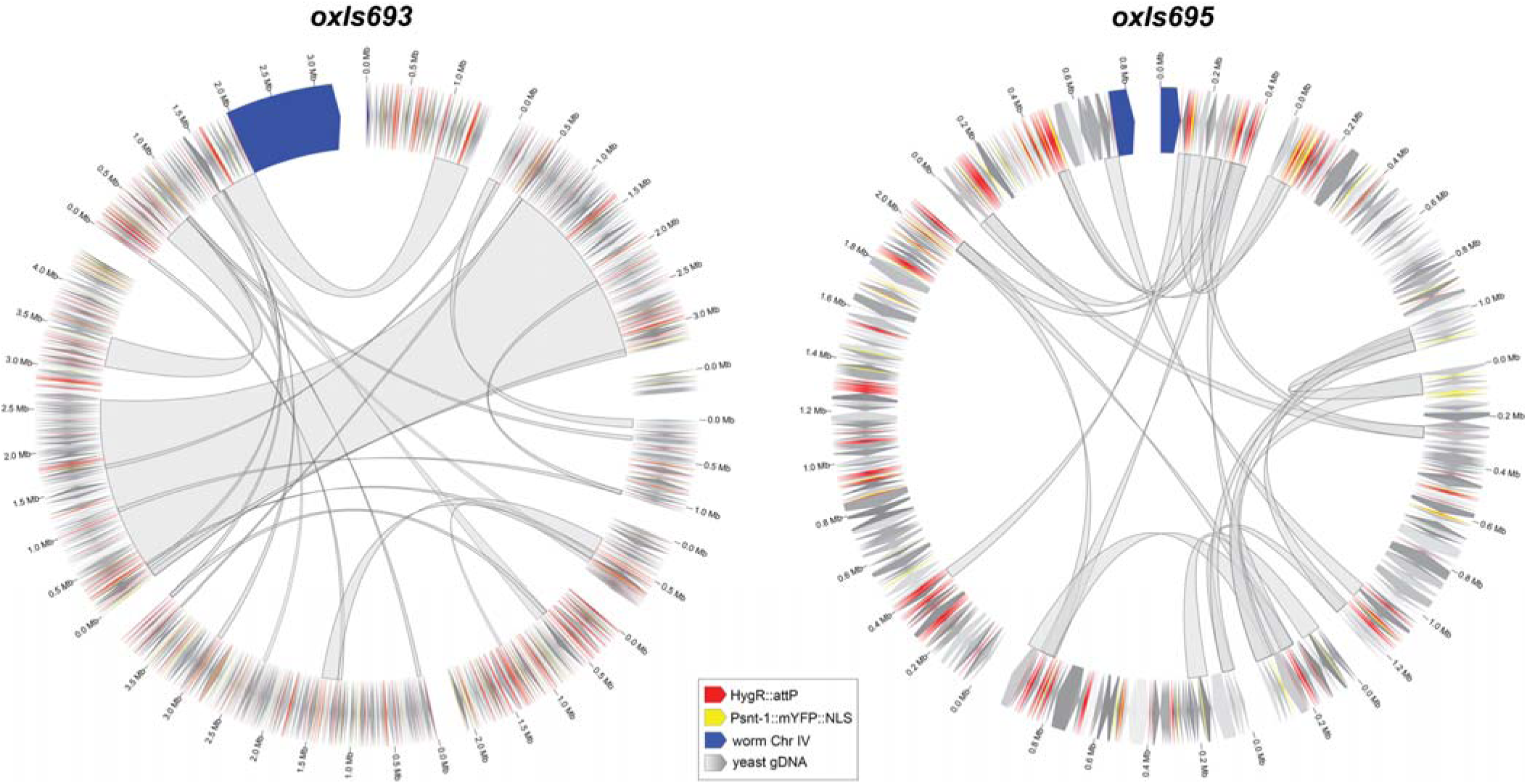
Large repeats found in integrated arrays. Repeats longer than 50 kb that are found on multiple contigs are plotted as gray arcs in Circos plots for *oxIs693* (left) and *oxIs695* (right). Around the outside of the plots, the annotated array contigs are plotted as in Figure 4. Gaps between contigs are caused by repeats that were too long for a sequencing read to span. Reads used for assembly were all at least 50 kb but no longer than 250 kb. Both arrays have 20 of these repeats.

**Figure S5.**
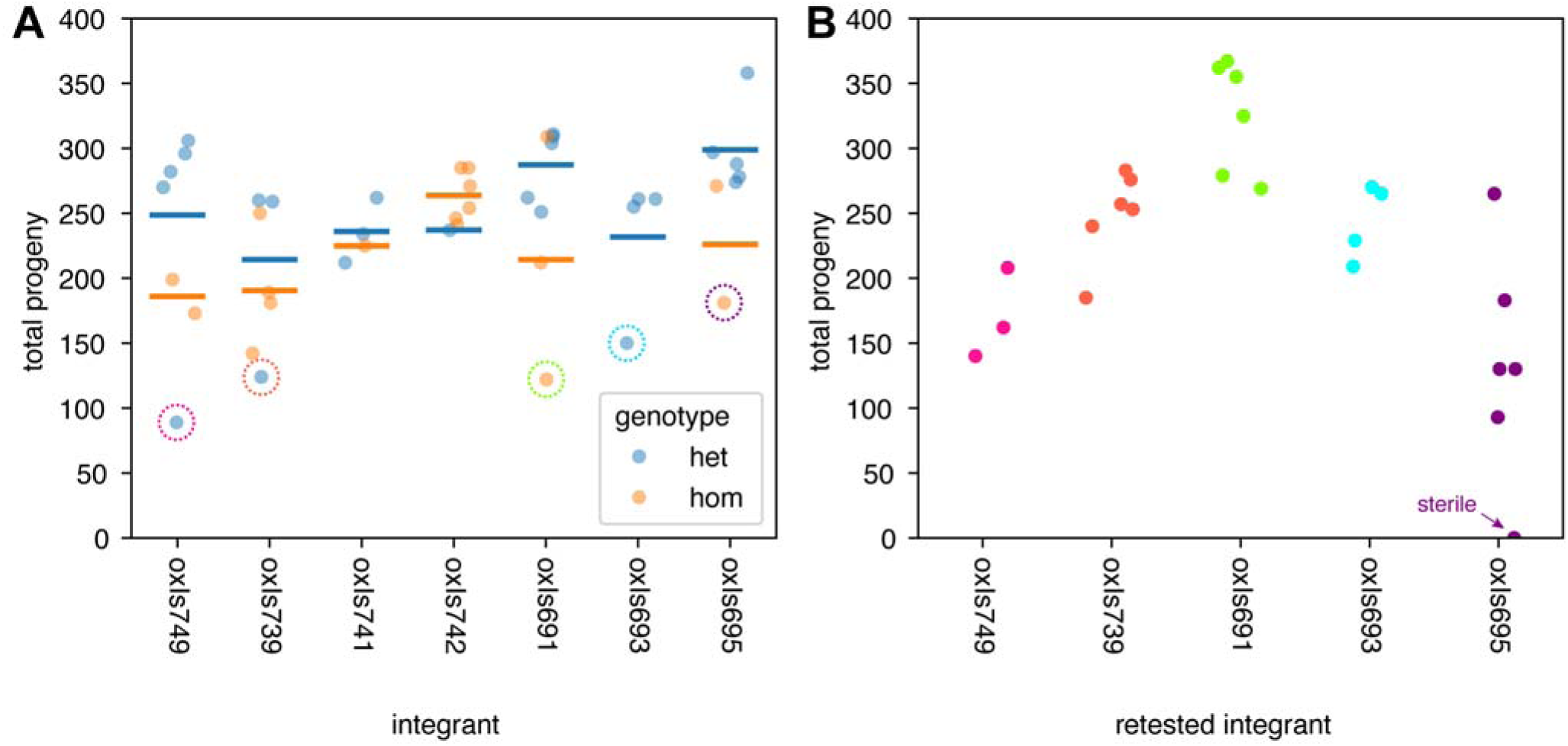
Fecundity of integrated array strains. (A) Array strains were outcrossed and their progeny counted. Orange: animals heterozygous for the array; Blue: animals homozygous for the array. Each point is the total progeny counted for a single animal, bars are means of the genotype. Circles indicate outliers with low fecundity. Progeny from these animals were retested in (B) (colored circles in A; correspond to colors in B). (B) Retesting fecundity in outliers. Progeny from circled animals in (A) were retested to determine whether their low-fecundity was hereditable. One retested line from *oxIs695* was sterile.

## Author Contributions

*Conceptualization* MSR, EMJ

*Funding Acquisition* EMJ, OR

*Investigation* MSR, RP, AH, YZ

*Methodology* MSR, RP

*Writing* MSR, EMJ

## Materials and Methods

### Reagents

All chemicals were purchased from Sigma-Aldrich (St. Louis, MO). Hygromycin, G418, and puromycin were purchased from GoldBio (St. Louis, MO). All enzymes were purchased from New England Biolabs (Beverly, MA). All synthetic oligonucleotide DNAs were purchased from Integrated DNA Technologies (Coralville, IA).

### C. elegans strains

*C. elegans* strains were cultured on nematode growth media (NGM) using standard methods (*39*). Worms were fed OP50 or HB101 bacteria. Animals were maintained at 15°C, 20°C, and 25°C. A list of worm strains created in this study can be found in Table S1. Drug selection plates were made by top-plating 500 ul of drug (hygromycin: 4 mg/ul, G418: 25 mg/ul, puromycin: 10 mg/ml + 0.1% Triton X-100) onto 6 cm NGM+HB101 plates.

### Molecular biology and cloning

A list of plasmids used in this study can be found in Table S2. A list of primers and guide RNAs used in this study can be found in Table S3. Novel plasmids were created using standard molecular biology techniques or ordered from Twist Bioscience. Annotated sequences for all plasmids can be found in File S2.

### General injection procedures

All plasmids were purified for injeciton using the Qiagen miniprep kit (Qiagen, Germantown, MD). Linear DNAs were prepared either by PCR using Q5 polymerase (New England Biolabs) or plasmid digestion followed by gel extraction using the Qiagen Minelute gel extraction kit (Qiagen). Details about injection mixes used here can be found in Table S4. All injections were conducted into day 1 or day 2 young adult hermaphrodites raised at 15°C on HB101 bacteria. In general, about 12-16 animals were injected in at least one gonad arm per injection session.

### Integration method

An injection mix containing transgene cargo, an *attP*-containing fragment/plasmid, a selection fragment/plasmid, and stuffer DNA was injected into young adult hermaphrodites that have an *attB* in their genome and express PhiC31 in their germline. Stuffer DNA was either a 1kb+ DNA ladder (Invitrogen, Carlsbad, CA, catalog #10787018) or yeast genomic DNA (Sigma-Aldrich catalog #69240-3). Injected animals were pooled 2-3 per NGM plate. After three days, array-bearing F1s were cloned to antibiotic selective plates (or NGM plates in the case of *unc-119* rescue) and grown at 25°C. When plates were close to starvation, a 1 cm x 1 cm chunk of agar was moved to a new selective plate. Plates can be chunked when close to starvation as many times as are necessary. In this manuscript, after two serial transfers strains were chunked to non-selective plates to grow animals for screening. The next day, 8 array-bearing L2-L4 animals per line were singled to non-selective plates. Three days later these plates were screened for homozygous array-bearing progeny. Results from all integration experiments can be found in File S1. A detailed protocol can be found in File S3.

### Integration without selection

EG10476 and EG10469 were crossed to generate an *oxSi1337[attB] I; oxSi1347[PhiC31] II* strain. These animals were injected with pSEM233 (a TagRFP selection marker at 1 ng/ul, linearized by NdeI digest), pMR791 (an *attP*-containing plasmid at 10 ng/ul, linearized by NotI digest) and Invitrogen 1kb+ DNA ladder (at 75 ng/ul). F1 array-bearing progeny were cloned to individual plates. About every three days, four animals from each line were cloned to new plates to create a set of four plates for each line. Plates from each line were screened for 100% array-bearing progeny at each generation.

### attB CRISPR and integration

EG10408 (*oxSi1347[PhiC31] II*) was injected with Cas9 (325 ng/ul final concentration, UC Berkeley QB3 Macrolab, Berkely CA), tracrRNA (100 pmol, IDT), an sgRNA for an insertion site (100 pmol, IDT), an ssODN for repairing with an attB site (100 pmol, IDT), and 100 ng/ul of array cargo (see Table S3 for crRNA and ssODN sequences). The selection and screening strategy was the same as for injection into a PhiC31 attB strain: array-bearing F1s were cloned to hygromycin plates and grown at 25C for cycles of starvation and chunking. Eight animals for each line were then singled and screened for homozygous integrated arrays.

### Calculating integration efficiency

Injected animals (P0 generation) were pooled 2-3 per plate after injection and allowed to lay eggs. F1 array-bearing animals were singled to new plates (containing drug if necessary for array selection, or NGM if using an *unc-119(+)* rescue). These animals were grown until either the plate starved or the line died because the array was not transmitted beyond the F1 generation. This set of lines (F2 array-bearing lines that survive the initial selection) make up the total set of lines and is the denominator of the efficiency calculation. The number of these lines that yields a homozygous integration is the numerator of the efficiency calculation. This calculation eliminates variability in array formation rate per injected animal, which can vary widely from many factors including injector skill, DNA mix quality, and injection strain health.

### Creation of oxC1

The recombination suppressor balancer *tmC20* balances the attB site on chromosome I. This balancer has an integrated array linked to *unc-14,* but has no phenotype. In order to add a recessive mutation with a visible phenotype to *tmC20,* we created a premature stop in *unc-40*. We injected EG10476 (*tmC20 I; oxSi1347[PhiC31] II*) with CRISPR reagents and fluorescent co-injection markers targeting residue 1138 in *unc-40* and screened the progeny of array-bearing F1s for *unc-40(null)* F2s. These were cloned and outcrossed to N2 to (1) isolate them from the PhiC31 transgene in the injection strain background, and (2) confirm that the *unc-40* mutation was linked to the Pmyo-2::YFP fluorescent marker present in the balancer.

### Creation of him-8(ox1602)

To create a fluorescently-marked Him chromosome, we used CRISPR to edit the *e1489* allele into EG7927, which carries *oxTi404*, a miniMos insertion of *Peft-3::TagRFP-T-NLS* approximately 0.2cM from him-8. We injected this strain with CRISPR reagents, an ssODN donor encoding *e1489*, and a pRF4 dominant roller plasmid, and screened the progeny of Rol F1s for Him F3s. These were outcrossed to N2 to (1) isolate from the *unc-119(ed3)* allele in the EG7927 background and (2) to confirm that RFP was linked with the Him phenotype.

### Fluorescent imaging

Confocal images were taken using a Zeiss LSM880 using a GaAsP detector. Epifluorescence images were taken on a Zeiss Axioskop compound fluorescence microscope equipped with a CoolLED (Andover, Hampshire, UK) pe-300lite white light LED and an Amscope (Irvine, CA) MF2003-BI color CMOS camera. Worms were anesthetized using 50 mM sodium azide before mounting on a 3% agarose pad for imaging. For images where direct comparisons were required, the same imaging settings were used for all samples.

### DNA extraction for whole-genome shotgun sequencing

*C. elegans* lines were grown on 10 cm NGM plates seeded with HB101 bacteria. Before starvation, worms were gently washed off plates with M9 and pelleted and washed 2x with M9 to remove bacteria. Genomic DNA was extracted using a Qiagen DNeasy Blood and Tissue kit (Qiagen). Whole genome sequencing was performed by the University of Utah Huntsman Cancer Center Genomics Core. Libraries were prepped using the New England Biolabs NEBNext Ultra II FS DNA Library Prep kit (catalog #E7805L) and sequenced on an Illumina NovaSeq X using a paired-end 2x151cycle protocol.

### Identifying array insertions from short read sequencing

FASTQ files were processed using a custom pipeline. Briefly, reads were mapped using bwa-mem (*40*) to a reference comprised of the *C. elegans* genome (WS276) and plasmid sequences contained in the injection mix of the sequenced strain (usually pSEM233 and pCFJ782). After alignment, structural variants were called with Manta (*41*), and filtered for variants that contained array transgenes using bcftools (*42*).

### Estimating array size from short read sequencing

Reads were mapped using bwa-mem (*40*) to a reference comprised of the *C. elegans* genome (WS276), the yeast genome (R64-sacCer3), and the sequences of plasmids pMR201 and pCFJ782. Array size was calculated based on the proportion of reads mapping to the worm genome vs mapping to the yeast genome and plasmids:

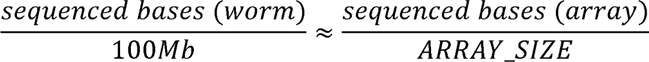

### Saturation analysis of yeast genomic DNA in arrays

All analyses were performed with bedtools (*56*) and custom python scripts (File S4). To measure saturation of the yeast genome in arrays, the yeast genome was binned into 1 kb fragments for all analyses, and short-read sequencing coverage of that bin was used as a proxy for incorporation of a yeast sequence into the array. To calculate mean saturation of a set of arrays, the order of the set of arrays was randomly permuted 1000 times and the mean cumulative yeast genome fraction after adding each array was calculated.

A Monte Carlo simulation was used to create a null distribution for the random assembly of arrays made from the yeast genome. For each simulated array, three parameters were chosen. (1) a total array size (in Mb) was drawn from a length distribution related to the estimated lengths of sequence arrays (*list(numpy.arange(1,5,0.1)) + [6,8,10,12,15,20]*). (2) A random fragment length was drawn from the distribution of lengths found in three arrays with long read sequences (*oxIs691, oxIs693, oxIs695*). (3) a random position in the yeast genome was chosen as the left end of that fragment. Fragments were accumulated in the “array” until the target size was reached. Twenty arrays were made per simulation iteration, and their mean saturation was calculated as for real arrays. The saturation null distribution was based on 1000 iterations of the simulation.

### DNA extraction for nanopore sequencing

*C. elegans* strains were grown to starvation on a 15cm 2XYT media plate that was seeded with NA22 bacteria. Starved worms were rinsed off the plate into a 15 mL conical with wash buffer (100mM Tris-HCl pH 8.0, 100mM NaCl, 20mM EDTA). To further remove any residual bacteria, worms were pelleted and washed three times with wash buffer, with the middle wash including a 30-minute incubation at room temperature to aid the removal of any gut bacteria. After pelleting, 50 µL of worms were resuspended in 485 µL of wash buffer and frozen at -80°C (*24*). The worms were subsequently thawed and Proteinase K and SDS were then added to final concentrations of 4 mg/mL and 0.2%, respectively. Samples were then incubated for 2 hours at 60°C. Following this initial digestion, 500 µL of lysis buffer (100mM Tris-HCl pH 8.0, 100mM NaCl, 20mM EDTA, 4mg/mL Proteinase K and 0.2% SDS) was added and the sample was incubated overnight. RNAse A was then added, and the sample was incubated at 30°C for 30 minutes. DNA was then extracted by performing a phenol-chloroform extraction and ethanol precipitation (*43–45*). DNA quality and quantity was checked using a NanoDrop spectrophotometer and a Thermo Fisher Scientific Qubit 4 Fluorometer.

Oxford Nanopore Technologies (ONT) libraries were prepared using ONT SQK-LSK114 according to refs (*24*, *44*). All samples were sequenced to over 50x coverage (>5Gb) using the ONT MinION sequencing device based on manufacturer’s instructions and previously published protocols (*24*, *44*). All basecalling was performed using the ONT MinKNOW software and the super-accurate basecalling model, with only passed reads and reads over 1kb being kept for downstream analysis. Sequencing quality was analyzed using NanoPlot (v.1.41.3) (*46*).

### Assembly of array reads

Integrated arrays were assembled by performing *de novo* genome assembly using Flye (v.2.9.1-b1780) (*47*) with the following parameters; --genome-size 120m --threads 56 --iterations 5 --min-overlap 10000--asm-coverage 40. Locations of the integrated arrays were confirmed by mapping the Flye contigs to the *C. elegans* reference genome (WBcel235) (*48*). The quality and completeness of each assembly was manually inspected using IGV (v.2.16.1) (*49*). When necessary, integrated array assemblies were corrected or extended using a combination of Winnowmap2 (v.2.03) (*50*) followed by Flye for polishing. More specifically, this meant mapping all reads to the integrated array contigs and checking for any soft-clipped reads in IGV that could be used to extend contigs. If the integrated array contigs could be extended, the sequences from the soft-clipped reads were then manually concatenated to the end of the contig and validated using Winnowmap2 and IGV. If correct, the new additions were polished using Flye. A similar method for elongating contigs was employed by (*51*). The sequences of the pCFJ782, pMR209, the *C. elegans* reference genome (WBcel235) (*48*), and the *S. cerevisiae* reference genome (R64-sacCer3) (*52*) were then mapped to the integrated array contigs using minimap2 (*53*). Estimates of array sizes from multi-contig assemblies was calculated as the total non-worm DNA contained within all contigs. As such, it is likely an overestimate of the actual size of the array, given that arrays are repetitive and there will be overlaps between contigs. Array contig sequences can be found in File S5.

### Analysis of array assemblies

All analyses were performed using custom Python scripts (File S4). Plots were created using matplotlib (*54*) and seaborn (*55*). To identify repeats within assemblies, contigs were aligned to each other using minimap2 (*54*) using default parameters. The resulting hits were filtered for those longer than 50 kb and with a mapping quality score greater than 50. Circos plots were drawn using pyCirclize (https://github.com/moshi4/pyCirclize).

