## Supplemental File 3 (Detailed Protocol) for "Targeted integration of extrachromosomal arrays in *C. elegans* using PhiC31 integrase"

**Annotated protocol for**

**PhiC31-mediated Integration of Arrays of Transgenes (PhIAT)**

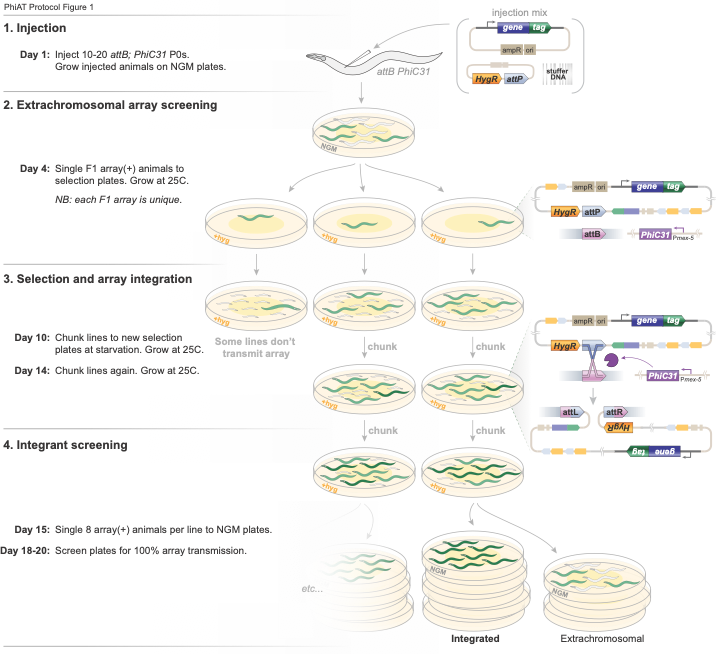

Rich *et al.* (2026) Targeted integration of extrachromosomal arrays in *C. elegans* using PhiC31 integrase

**PhIAT protocol** ^1^

**0.** **Creating injection strain** ^2^

*Day -10*	Cross *attB; RFP Him* males to *balancer; PhiC31::mNG* hermaphrodites ^3^.

*Day -7*	Single *attB/balancer; PhiC31::mNG/+; RFP Him/+* hermaphrodite F1 cross-progeny.

*Day -4*	Single 8x *balancer(-)* *RFP(-) PhiC31::mNG(+)* hermaphrodites to HB101 plates. Grow at 15C.

*Day -1*	Screen adults for homozygous *PhiC31::mNG(+)* lines. Keep one line at 15C ^4^.

**1. Injection**

*Day 0*	Grow *attB PhiC31* strain on HB101 plates at 15C ^4^.

*Day 1*	Make injection mix that includes transgenes, selection markers, and *attP*. See note (5) below for details.

Inject 10-20 animals ^6^.

Place injected animals on HB101 plates. Incubate at 25C.

**2. Extrachromosomal array screening**

*Day 4*	Single F1 array(+) animals to new HB101 plates, supplemented with drug selection if necessary ^7^. Incubate plates at 25C.

*Note: all F1 arrays are unique. Single many to get diverse integrants.*

**3. Selection**

*Day 10*	Chunk surviving lines to new selection plates ^8^.

*Day 14*	Chunk surviving lines to new selection plates.

**4. Screening** ^9^

*Day 15*	Single 8 array(+) animals per line to **NGM** plates ^10^.

*Day 18-20*	Screen plates for 100% array transmission. These are integrants ^11^.

**5. Outcross**

*Day 21*	Cross integrant to N2 ^12^ males to outcross and remove PhiC31.

**PhIAT protocol notes:**

1	See PhIAT protocol illustrated steps (Protocol Fig. 1)

2	PhiC31 will cut an *attB* site without an *attP* present, eventually mutating the *attB* and eliminating integration. As such, for long-term storage, we maintain *attB* and *PhiC31* strains separately. Before integration injections, the strains are crossed to reconstitute an injectable integration strain.

3	The strains you cross depends on which *attB* you want to integrate at. To simplify reconstitution crosses, we made PhiC31 strains containing *attB* balancers and *attB* strains containing fluorescently-marked Him alleles:

| attB strain | x | balancer + PhiC31 strain |
| --- | --- | --- |
| EG10541 *attB I; RFP him-8 IV* | x | EG10478 *Bal[unc-40(-), YFP] I; PhiC31[mNG] II* |
| EG10542 *attB II; RFP him-8 IV* | x | EG10570 *Bal[dpy-10(-) unc-52(-), GFP] II / mnDf61[lethal]; PhiC31[mNG] IV* |
| EG10543 *attB IV; him-5 RFP V* | x | EG10473 *PhiC31[mNG] II; Bal[unc-5(-), YFP] IV* |

Table 1. *attB*s and corresponding PhiC31+balancers for reconstituting injection strains.

For example, to make an injection strain for the *attB* site on chromosome IV, you would follow the cross shown in Protocol Fig. 2.

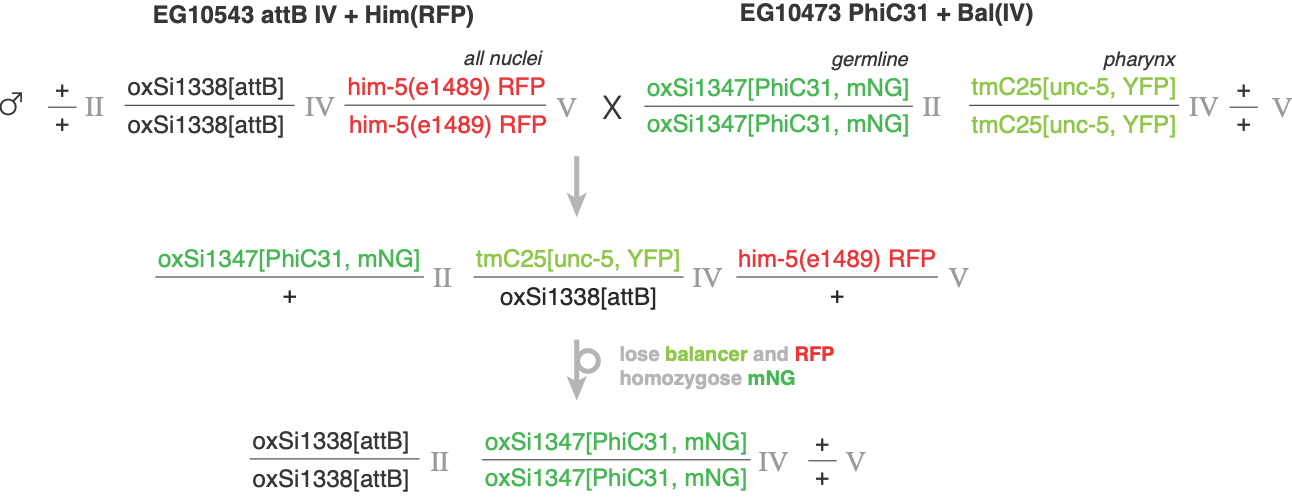

Protocol Figure 2. Example injection strain reconstitution cross.

Heterozygotes can also be injected and will yield integrants. In this method, we recommend pooling injected animals and screening the whole pool for integrants.

4	*attB* mutagenesis occurs at a non-trivial (but currently unknown) rate in the germline of an *attB PhiC31* strain. As such, we recommend propagating injection strains only when necessary to limit the number of generations they go through (and therefore limit the frequency of mutated *attB* sites in the population).

5	A PhIAT injection has 4 components: your transgenes, a selectable marker, *attP* sites, and a fluorescent array marker. Arrays can be made either with linear DNAs (in which *attP* sites are appended by PCR to selection plasmids) or plasmid DNAs (in which pMR903 is coinjected along with transgene and selection plasmids). Note: we usually find that expression from linear arrays is much stronger than in plasmid arrays, and so the relative amount of transgene plasmids should be reduced, sometimes over ten-fold. See Tables 2 and 3 for examples; these injections yielded similarly-bright arrays.

| DNA | ng/ul |
| --- | --- |
| pMR903 (attP) | 6 |
| pSEM233 (Pmlc-1::TagRFP) | 11 |
| pCFJ782 (HygR) | 10 |
| DNA ladder | 75 |

Table 3. Plasmid PhIAT injection

| DNA | ng/ul |
| --- | --- |
| HygR::attP PCR (from pCFJ782) | 2.5 |
| Pmlc-1::TagRFP PCR (from pSEM233) | 1 |
| DNA ladder | 100 |

Table 2. Linear PhIAT injection

6	The number of injected animals may vary depending on injector skill. We aim for having ~50 F1 arrays, and routinely get this many arrays by injecting about 10 animals in a single gonad arm.

7	We have integrated arrays using hygromycin B, neomycin/G418, and puromycin drug selections, as well as *unc-119(+)* rescue (see Table 4 for drug concentrations and resistance plasmids).

For drug selections, top-plate 500ul of drug stock on a 4cm plate. Let the plate dry with the lid off (this usually takes about 30-60 minutes) before storage. We purchase our drugs from Gold Bio.

8	In our hands, 30-50% of F1 arrays are transmitted to the next generation.

9	The primary method we use to isolate integrants is screening for homozygous segregation of a fluorescent marker. Integrants can also be identified by PCR, as the *attB+attP -> attL+attR* recombination is predictable (unpublished). ApE has a recombination tool that allows for testing these recombinations *in silico* and designing primers to genotype them (<https://jorgensen.biology.utah.edu/wayned/ape/>).

10	The number of animals to single for screening is arbitrary. We recommend eight for two reasons: first, it’s a large enough set to not miss relatively low-frequency integrants, and second, a stack of eight plates fits well in our incubators. Integrants will be isolated from using fewer plates per line, but some might be missed. For screening large numbers of lines, we have used GelDrops (PMID: 40985011) on the lids of 96-well plates.

| Selection | Stock conc. | Plasmid |
| --- | --- | --- |
| Hygromycin B | 4 mg/ml | pCFJ782 |
| Neomycin/G418 | 25 mg/ml | pCFJ594 |
| Puromycin | 10 mg/ml + 0.1% Triton100 | pCFJ744 |
| *unc-119(-)* | n/a | pMR201 |

Table 4. Selections and their plasmids.

11	Because PhIAT uses a selection, any array that transmits near 100% will increase in population frequency during selection. The primary form of false-positives during screening are extrachromosomal arrays that transmit near 100%. To get rid of these, we grow putative integrant plates for 2-3 generations after singling; after two generations a plate with a high-transmission array will have many array(-) animals, while a true integrant will have zero.

12	We recommend outcrossing integrants to remove the PhiC31 transgene and limit off-target mutations in the strain background. While this cross can be done with a wild-type strain, we recommend using a visible mapping marker to confirm the array inserted at the correct site. Some good mapping markers are:

- *oxSi1337* I@-5.3 vs *dpy-5(e61)* I@0
- *oxSi1270* II@0.8 vs *dpy-10(e128)* II@0
- *oxSi1338* IV@1.4 vs *unc-22(e66)*

**Strains list** (see Table S1 for full list of strains.)

| *PhiC31 strains* |  |  |  |
| --- | --- | --- | --- |
| strain | **description** | **genotype** | |
| EG10408 | PhiC31 II | oxSi1347[Pmex-5::PhiC31(PATC)::sl2::mNeongreen::glh-2UTR *jsSi1579] II | |
| NM5406 | PhiC31 IV | jsSi1623[loxP, Pmex-5::phiC31::SL2::mNeongreen::glh-2UTR, FRT3] IV | |
| EG10476 | PhiC31 with balancer for attB I | tmC20[unc-14(tmIs1219)] I ; oxSi1347[Pmex-5::PhiC31(PATC)::sl2::mNeongreen::glh-2UTR, *jsSi1579] II | |
| EG10570 | PhiC31 with balancer for attB II | mnC1[dpy-10(e128) umnIs32(Pmyo-2::GFP) unc-52(e444)] / mnDf61[lethal] II ;  jsSi1623[loxP Pmex 5::PhiC31::SL2::mNeonGreen::glh-2UTR FRT3] IV | |
| EG10473 | PhiC31 with balancer for attB IV | oxSi1347[Pmex-5::PhiC31(PATC)::sl2::mNeongreen::glh-2UTR, *jsSi1579] II ;  tmC25 [unc-5(tmIs1241[Pmyo-2::YFP])] IV | |
| *attB strains* |  |  |  |
| strain | **description** | | **genotype** |
| EG10469 | attB near ttTi4348 I | | oxSi1337[attB, *ttTi4348] I |
| EG10470 | attB near ttTi5605 II | | oxSi1270[attB, *ttTi5605] II |
| EG10471 | attB near cxTi10816 IV | | oxSi1338[attB, *cxTi10816] IV |
| EG10541 | attB I + RFP him-8 IV | | oxSi1337[attB] I ; oxTi404[Peft-3::tdTomato::H2B::unc-54 cb-unc-119(+)] him-8(ox1602) IV |
| EG10542 | attB II + RFP him-8 IV | | oxSi1270[attB] II ; oxTi404[Peft-3::tdTomato::H2B::unc-54 cb-unc-119(+)] him-8(ox1602) IV |
| EG10543 | attB IV + him-5 RFP V | | oxSi1338[attB] IV ; him-5(e1490) oxTi405[Peft-3::tdTomato::H2B::unc-54 cb-unc-119(+)] V |
| EG10481 | CyOFP(loxP-attB-loxP) I | | oxSi1411[Pelo-5::CyOFP(loxP-attB-loxP), *oxTi185] I ; unc-119(ed3) III |
| EG10567 | CyOFP(attB) I | | oxSi1451[Pelo-5::CyOFP(attB)::rps-1UTR, *jsSi1727] I |
| EG10568 | CyOFP(attB) II | | oxSi1452[Pelo-5::CyOFP(attB)::rps-1UTR, *jsSi1726] II |
| EG10569 | CyOFP(attB) IV | | oxSi1453[Pelo-5::CyOFP(attB)::rps-1UTR, *jsSi1986] IV |

**Plasmids and primers lists** (see Table S2 for full list of plasmids and primers)

| *attP and selection plasmids* | |
| --- | --- |
| plasmid | **description** |
| pMR903 | 197bp attP |
| pMR791 | 39bp attP (low efficiency) |
| pCFJ594 | G418R |
| pCFJ707 | PuroR |
| pCFJ782 | HygR |
| pMR201 | Cbr-unc-119(+) |

| *fluorescent plasmids used in manuscript* | |
| --- | --- |
| plasmid | **description** |
| pSEM233 | Pmlc-1::TagRFP-T::tbb-2UTR (pan-muscle, cytosolic) |
| pMR196 | Psnt-1::YFP::histone::unc-54UTR (pan-neuronal, nuclear localized) |

| primers | sequence (5'-3') | description |
| --- | --- | --- |
| M13_F | TGTAAAACGACGGCCAGT |  |
| attP::M13R | CGCCCCCAACTGAGAGAACTCAAAGGTTACCCCAGTTGGGGcaggaaacagctatgaccatg | attP (capitalized) appended to M13R |
